# Synphilin-1 mitigates autophagy dysfunction, modulates ubiquitinated protein aggregation, and promotes cell survival during proteotoxic stress

**DOI:** 10.64898/2026.05.31.729136

**Authors:** Nadine M Lebek, Kenneth G Campellone

## Abstract

The decline of cellular proteostasis is a hallmark of aging and key contributor to neurodegenerative diseases. Protein turnover is controlled by the ubiquitin-proteasome and autophagosome-lysosome systems, but how degradation is coordinated when one of these pathways is compromised is not well understood. To study the regulation of proteostasis, we utilized human fibroblasts with targeted knockouts of the cytoskeletal factors WHAMM and JMY, which control multiple steps in autophagy. We found that cells lacking both WHAMM and JMY accumulated numerous intense foci of ubiquitinated proteins when exposed to proteotoxic stress and relied on proteasomes to clear the foci when the stressor was removed. RNA-seq and immunoblotting revealed that WHAMM/JMY knockout cells increased their expression of Synphilin-1, an α-synuclein-interacting protein implicated in Parkinson’s Disease. In WHAMM/JMY knockout cells that upregulated endogenous Synphilin-1, and in cell lines engineered to overexpress mCherry-Synphilin-1, ubiquitinated proteins were present in structures containing both Synphilin-1 and proteasomes. RNAi-mediated depletion of Synphilin-1 caused a buildup of ubiquitinated proteins and the ubiquitin-binding adaptor protein SQSTM1/p62, while decreasing cell survival in response to proteotoxic stress. These data suggest that Synphilin-1 plays a pro-survival role in cells with impaired autophagy and functions in the distribution of ubiquitinated cargo during proteasomal degradation.

## INTRODUCTION

Proteostasis, or protein homeostasis, is achieved within cells through the coordinated production, folding, modification, trafficking, and removal of proteins. The maintenance of a functional proteome allows cells to proliferate and respond to diverse stimuli, whereas alterations in biosynthetic and degradative processes are linked to impaired stress responses and increased disease pathologies (Labbadia and Morimoto, 2015; Yerbury *et al*., 2016; Wilson *et al*., 2023). In particular, defects in the protein degradation machinery and its regulators can lead to accelerated biological aging and protein aggregation disorders (Kaushik and Cuervo, 2015; Hipp *et al*., 2019). However, the mechanisms that maintain proteostasis when some degradation routes are compromised are not well understood.

Protein degradation in mammalian cells is controlled by the ubiquitin-proteasome system (UPS) and the autophagosome-lysosome system (ALS) (Dikic, 2017; Ji and Kwon, 2017; Aman *et al*., 2021). Both systems involve the conjugation of ubiquitin to lysine residues in proteins, thus targeting them to degradative pathways (Kwon and Ciechanover, 2017; Yin *et al*., 2020). In UPS-mediated degradation, misfolded proteins and other polyubiquitinated substrates are engaged by proteasomes where they are unfolded by the regulatory proteasomal subunit and fed through the proteolytic core (Livneh *et al*., 2016; Collins and Goldberg, 2017). Larger substrates, such as organelles and some protein aggregates, that are unable to be unfolded for proteolysis via the proteasome are processed by the ALS. In selective forms of macroautophagy (hereafter referred to as autophagy), ubiquitinated substrates are bound by autophagy adaptors/receptors, which additionally interact with lipidated LC3-family proteins present in nascent autophagosomal membranes (Lamark and Johansen, 2021; Vargas *et al*., 2023). Mature double membrane-bound autophagosomes then fuse with lysosomes for digestion of their macromolecular content (Nakamura and Yoshimori, 2017; Zhao and Zhang, 2019). While the two-pronged UPS/ALS approach to intracellular degradation effectively limits the presence of unwanted proteins, each system can still become overwhelmed or inhibited by proteotoxic stressors that create an overabundance of substrates.

Misfolded proteins have the potential to form small, highly toxic oligomers or larger, possibly less damaging, aggregates. The sequestration of proteins into aggregates, inclusion bodies, phase-separated condensates, or higher-order assemblies called aggresomes can therefore be considered a third proteostatic approach for managing harmful protein species (Sontag *et al*., 2017; Amzallag and Hornstein, 2022; Hipp and Hartl, 2024). The biogenesis of such structures acts as a cytoprotective mechanism until the UPS and ALS regain their degradative capacities (Kawaguchi *et al*., 2003). Chaperone-based disaggregation processes can extract proteins from aggregates and refold them or allow them to be degraded through the UPS (Mogk *et al*., 2018). Alternatively, aggregation and targeting to selective autophagy pathways can be modulated through autophagy adaptors/receptors such as SQSTM1/p62 (Pankiv *et al*., 2007; Kirkin *et al*., 2009; Sarraf *et al*., 2020; Bauer *et al*., 2023).

The burden of misfolded proteins increases with age, and the UPS and ALS pathways become less efficient, leading to a progressive decline in proteostasis in cells from mice and humans (Carnio *et al*., 2014; Leeman *et al*., 2018; Sabath *et al*., 2020). The UPS and ALS are also acutely impaired by the production of aggregation-prone proteins such as α-synuclein and huntingtin, and dysfunctional proteostasis is therefore a major contributor to neurodegenerative diseases like Parkinson’s and Huntington’s (Wyttenbach *et al*., 2000; Cuervo *et al*., 2004; Emmanouilidou *et al*., 2010; Winslow *et al*., 2010). In some circumstances, cellular upregulation of the ALS can partially compensate for a compromised UPS (Pandey *et al*., 2007; Zhu *et al*., 2010), and UPS activation may take place in response to ALS inhibition (Tannous *et al*., 2008; Wang *et al*., 2013), but UPS activity may also be suppressed by prolonged ALS disruption (Korolchuk *et al*., 2009).

These protein management systems are regulated by diverse factors, and in recent years the actin cytoskeleton has both emerged as a significant player in autophagy and been implicated in aggregation/disaggregation processes (Campellone *et al*., 2023). Actin monomers are assembled into filaments by proteins called nucleators, including the Arp2/3 complex, which cooperates with nucleation-promoting factors from the Wiskott-Aldrich Syndrome Protein (WASP) family to create branched actin filaments (Kramer *et al*., 2022). While Arp2/3 complex activity may control the disaggregation of ubiquitinated mitochondria and the dynamics of p62 bodies (Hsieh and Yang, 2019; Feng *et al*., 2022), the functions of Arp2/3 and WASP-family members are better understood during autophagy.

The founding mammalian Arp2/3 activator, WASP, affects autophagic processes specifically in hematopoietic cells (Lee *et al*., 2017; Rivers *et al*., 2020), and the ubiquitously-expressed nucleation factors Cortactin, WASH, WHAMM, and JMY operate throughout the canonical autophagy pathway in diverse cell types. WHAMM and JMY activate the Arp2/3 complex during membrane remodeling processes ranging from autophagosome biogenesis and maturation (Coutts and La Thangue, 2015; Kast *et al*., 2015; Mathiowetz *et al*., 2017), to cargo receptor capture during autophagosome closure (Coulter *et al*., 2024), autophagosome rocketing (Kast *et al*., 2015; Hu and Mullins, 2019), autolysosome tubulation (Dai *et al*., 2019; Wu *et al*., 2021), and lysosomal repair (Theodore *et al*., 2026).

Despite the large body of work characterizing the functions of the nucleation factors in various steps of organelle dynamics, much less is understood about the impact of these proteins on overall proteostasis. How cells lacking the cytoskeletal modulators of autophagy are able to mitigate endogenous or acute proteotoxic stress is also unclear. In the current study, we used WHAMM/JMY double knockout (DKO) human fibroblasts to determine how nucleation factor-deficient cells coordinate the activities of the UPS, ALS, and protein aggregation-disaggregation systems in response to stress. We found that the α-synuclein-interacting protein Synphilin-1 is upregulated in WHAMM/JMY^DKO^ cells, where it functions as a central player in proteostasis by influencing the organization of ubiquitinated proteins, proteasomes, and proteotoxic stress responses.

## RESULTS

### Deletion of WHAMM and JMY decreases lipidation of the autophagosomal protein LC3

Defects in autophagosome biogenesis, movement, and tubulation have been found in cells lacking WHAMM or JMY individually (Coutts and La Thangue, 2015; Kast *et al*., 2015; Mathiowetz *et al*., 2017; Dai *et al*., 2019; Coulter *et al*., 2024). However, no autophagy-related phenotypes have yet been examined in cellular contexts where both WHAMM and JMY are absent. To explore the combined effect of WHAMM and JMY deficiency on autophagy, we utilized a WHAMM/JMY double knockout cell line derived from the parental human fibroblast-like eHAP cell line (King *et al*., 2021). Similar to previous experiments (Mathiowetz *et al*., 2017), we treated eHAP and knockout cells with chloroquine (CQ) for up to 2h to block lysosome-mediated degradation of autophagosomes, and immunoblotted cell extracts with antibodies that recognize proteins from the LC3 family, which exist in immature (LC3-I) and autophagosome-associated lipidated (LC3-II) forms (Klionsky *et al*., 2021).

Before the addition of chloroquine, eHAP parental cells and WHAMM/JMY^DKO^ cells contained similarly low levels of LC3-II, but the DKO cells harbored greater amounts of LC3-I, normalized to tubulin and GAPDH loading controls (Figure 1A). Upon exposure to chloroquine, parental cells showed a noticeable increase in LC3-II levels over time, whereas in DKO cells LC3-II was comparably less abundant (Figure 1A). Quantification of the LC3 species showed that basal LC3-I levels were nearly 4-fold higher and chloroquine-driven LC3-II accumulation was more than 4-fold lower in DKO cells (Figure 1B). Consistent with the latter results, the LC3-II:I ratio under chloroquine conditions was significantly reduced in DKO cells compared to parental cells (Supplemental Figure S1). The abundance and lipidation of GABARAP-family proteins, which have similar properties to the LC3s but play distinct roles in the progression of autophagy, did not differ between the parental and DKO cells (Supplemental Figure S1). These results suggest that WHAMM/JMY^DKO^ cells have a specific deficiency in converting LC3-I to LC3-II.

**Figure 1.**
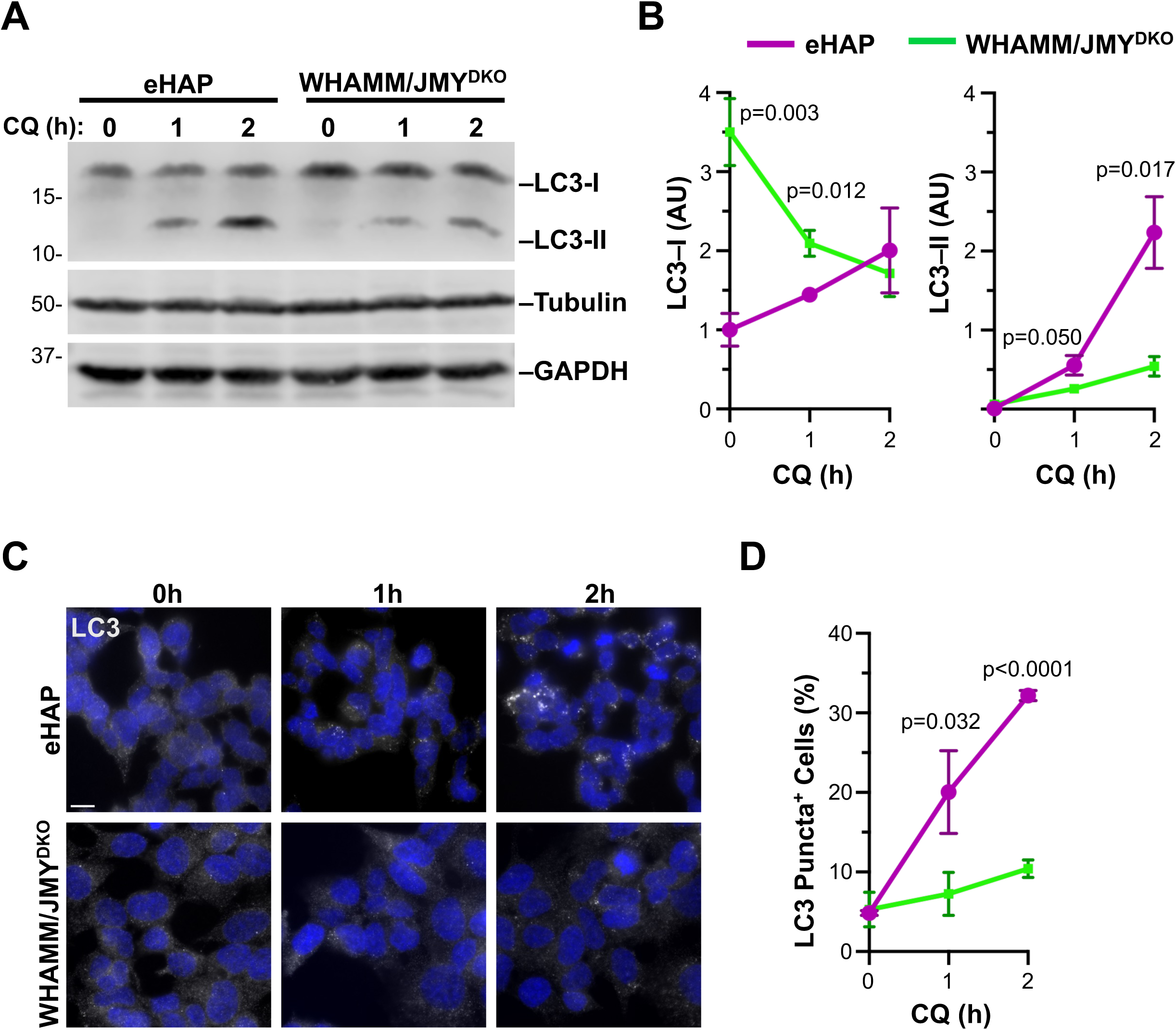
Inactivation of both WHAMM and JMY reduces lipidation of LC3. **(A)** eHAP and WHAMM/JMY^DKO^ cells were treated with chloroquine (CQ) for 0,1, or 2h, then lysed and subjected to SDS-PAGE and immunoblotted with antibodies to LC3, tubulin, and GAPDH. **(B)** LC3-I and LC3-II intensities from (A) were measured relative to tubulin and GAPDH and normalized to eHAP at 0h. Each point represents the mean from n=3 experiments ± SD. Significant *p* values comparing eHAP to DKO at each timepoint are noted (unpaired *t* tests). **(C)** eHAP and WHAMM/JMY^DKO^ cells were treated with CQ for 0, 1, or 2h and stained with an antibody to visualize LC3 (white) and DAPI to visualize DNA (blue). Scale bar: 10μm. **(D)** The number of cells with LC3 puncta were counted from (C) and expressed as a percentage of total cells counted (170-419 cells per condition per experiment). Each point represents the mean from n=3 experiments ± SD.

To visualize autophagosomal structures in eHAP and WHAMM/JMY^DKO^ cells, we next stained the cells with LC3 antibodies and examined them by fluorescence microscopy. Using this approach, LC3-II-associated membranes generally appear as bright puncta. In accordance with the low steady-state levels of LC3-II that were detected via immunoblotting, LC3 puncta were rarely observed in either parental or DKO cells in the absence of chloroquine (Figure 1C). Upon chloroquine treatment, however, approximately 20% of eHAP cells contained LC3 puncta by 1h, and this frequency rose to more than 30% by 2h (Figure 1, C and D). In contrast, the proportion of DKO cells harboring LC3 puncta barely reached 10% by 2h (Figure 1, C and D). Taken together, these immunoblotting and immunofluorescence data indicate that a combined deletion of WHAMM and JMY causes defects in LC3 lipidation.

### Ubiquitinated proteins accumulate in WHAMM/JMY^DKO^ cells following proteotoxic stress

Since autophagosome formation was impaired in WHAMM/JMY-deficient cells, we sought to evaluate how these cells respond to proteotoxic stress. We therefore treated parental and WHAMM/JMY^DKO^ cells with the antibiotic puromycin, which incorporates into nascent polypeptide chains, forcing their premature release from the ribosome and creating an abundance of misfolded polypeptides (Eggers *et al*., 1997). To first ask if puromycin exposure would impact cell viability, we compared the sensitivity of eHAP and WHAMM/JMY^DKO^ cells to a low dose of puromycin (0.25μg/mL) over an extended period of time (24h). Propidium iodide (PI) was added to the culture media to stain nucleic acids in dying/dead cells before fixing the cells and staining them with DAPI to visualize all nuclei. While <0.5% of cells were PI-positive when grown in normal media, approximately 2% of eHAP and nearly 4% of WHAMM/JMY^DKO^ cells were PI-positive following puromycin treatment (Supplemental Figure S2).

To explore the impact of a more acute proteotoxic stress, we treated each set of cells with a higher dose of puromycin (2μg/mL) and assessed the number of remaining adherent cells after 24h by counting the number of DAPI-stained nuclei per field of view. Under these conditions, about 4% of eHAP cells but only ∼0.75% of WHAMM/JMY^DKO^ cells remained (Supplemental Figure S2). The DKO cells were also more sensitive than eHAP cells to other ribosome-targeting drugs including geneticin and hygromycin (see Materials and Methods). Collectively, these observations show that compared to wild type cells, WHAMM/JMY-deficient cells are more likely to lose viability when exposed to proteotoxic stressors.

To examine the intracellular responses to acute puromycin treatment, we next subjected eHAP and WHAMM/JMY^DKO^ cells to the high dose of puromycin for a shorter period of time (3h), then fixed and stained the cells with an antibody to ubiquitin-conjugated proteins. In normal media, both eHAP and WHAMM/JMY^DKO^ cells showed punctate ubiquitin staining that was primarily restricted to the nucleus (Figure 2A). After puromycin treatment, both eHAP and WHAMM/JMY^DKO^ cells formed distinct ubiquitin foci in the cytosol that were larger and dramatically brighter than the nuclear ubiquitin puncta seen at steady state (Figure 2A). Quantification of the area of ubiquitin staining per puromycin-treated cell revealed that ubiquitin staining occupied a nearly 30% larger area within WHAMM/JMY^DKO^ cells than in parental cells (Figure 2B). The intensity of the ubiquitin staining was also 2-fold greater in WHAMM/JMY^DKO^ cells compared to eHAP cells (Figure 2C), likely reflecting a higher density of ubiquitinated cargo within the foci. These results suggest that the greater susceptibility of WHAMM/JMY-deficient cells to proteotoxic stress is related to an inability to efficiently degrade misfolded proteins.

**Figure 2.**
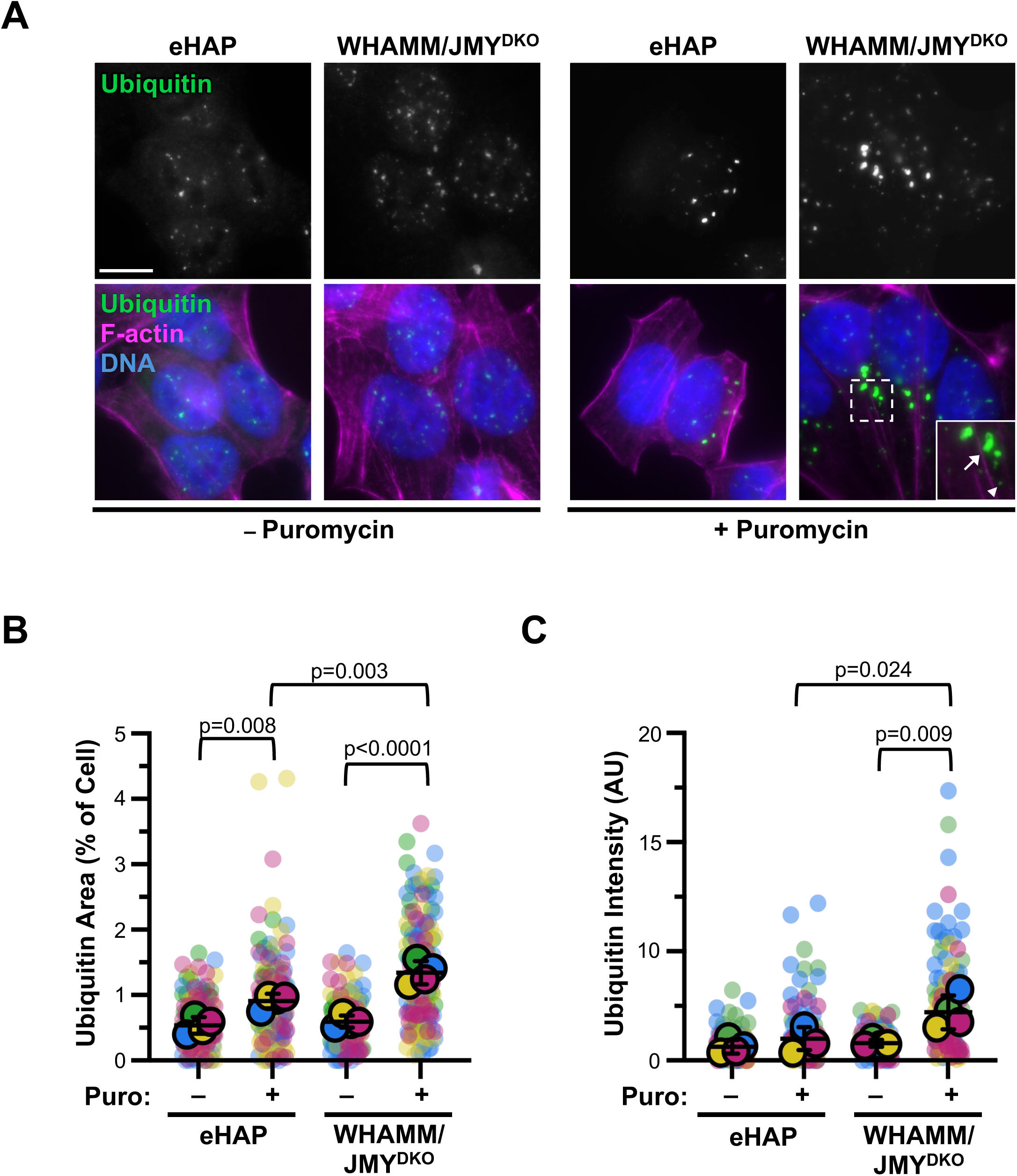
Proteotoxic stress causes WHAMM/JMY^DKO^ cells to accumulate ubiquitin foci. **(A)** eHAP and WHAMM/JMY^DKO^ cells were treated with 2μg/mL puromycin for 3h and stained with an antibody to visualize ubiquitin-conjugated proteins (green), phalloidin to visualize F-actin (magenta), and DAPI to visualize DNA (blue). The magnified insert shows examples of ubiquitin foci (arrow) and ubiquitin puncta (arrowhead). Scale bar: 10μm. **(B)** The area occupied by ubiquitin structures was quantified from (A). Each small symbol represents a single cell (23-40 cells per experiment), each larger symbol represents the average from n=4 experiments ± SD. Each experiment is coordinated by color. Significant *p* values are noted (ANOVA, Tukey post-hoc tests). **(C)** The intensity of ubiquitin staining in puromycin-treated cells was measured and normalized to the eHAP average and presented as in panel B.

### WHAMM/JMY^DKO^ cells rely on proteasomes to remove ubiquitin foci

Based on the puromycin-driven increase of ubiquitin foci in WHAMM/JMY^DKO^ cells, we next examined whether the cells were capable of removing these structures after taking away the proteotoxic stressor. We therefore treated eHAP and WHAMM/JMY^DKO^ cells with normal control media or media containing puromycin for 3h, then washed out the drug-containing media and replaced it with drug-free media for an additional 3h to allow cells to recover before fixing and staining them for ubiquitin. While both the area and the intensity of ubiquitin staining was higher in the WHAMM/JMY^DKO^ cells than in the eHAP cells after 3h of puromycin treatment, ubiquitinated protein levels in both cell lines returned to near-steady state levels after washout and recovery (Figure 3, A and B). This suggests that the WHAMM/JMY-deficient cells are still capable of removing ubiquitinated proteins.

**Figure 3.**
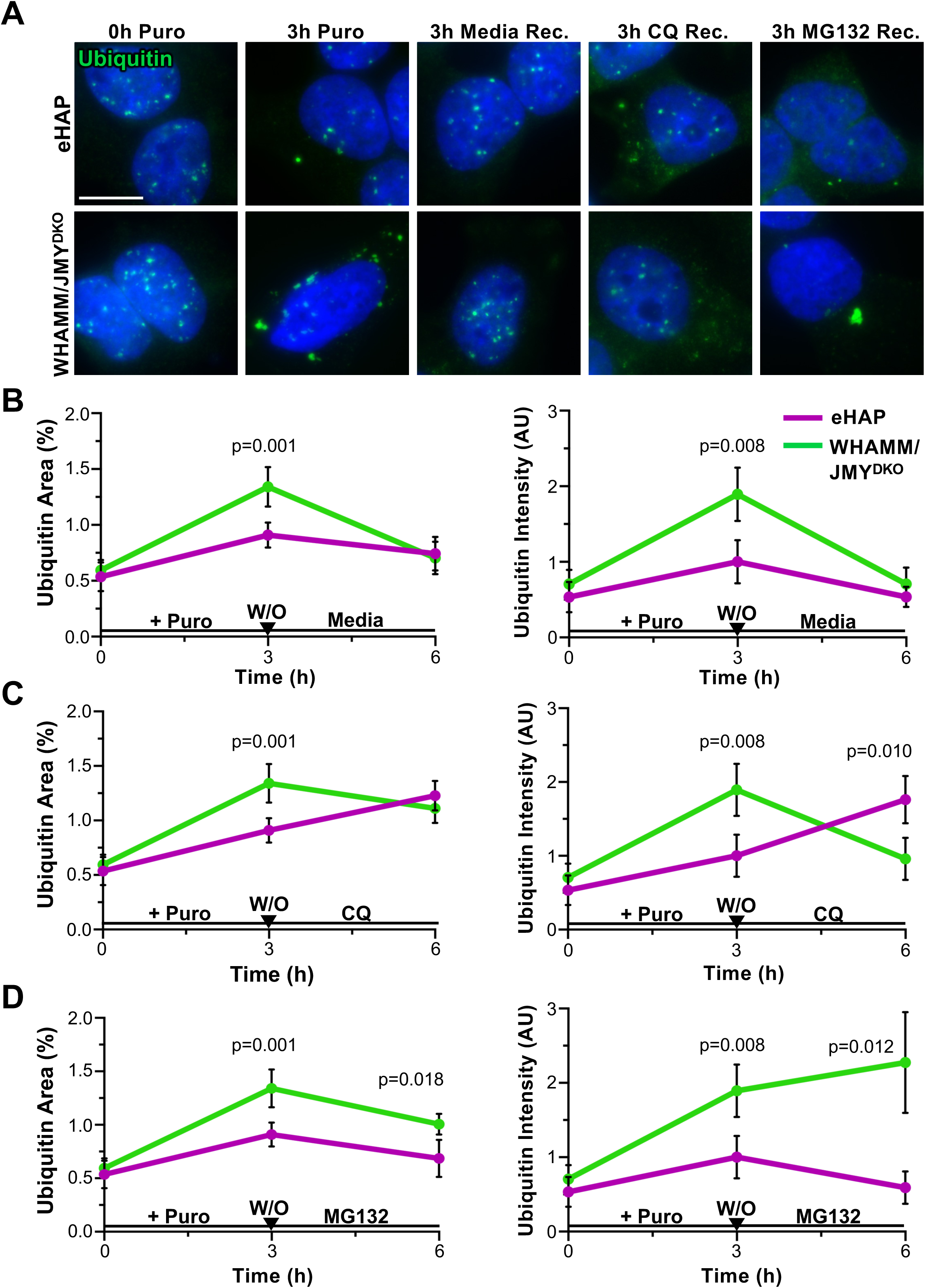
The proteasome is necessary for clearing ubiquitin foci in WHAMM/JMY^DKO^ cells. **(A)** eHAP and WHAMM/JMY^DKO^ cells were treated with 2μg/mL puromycin for 3h then subjected to a media washout (W/O) and allowed to recover (Rec.) for 3h in media, media + CQ, or media + MG132. Cells were fixed at 0h (untreated), 3h, and 6h timepoints and stained with an antibody to ubiquitin-conjugated proteins (green) and DAPI to visualize DNA (blue). Scale bar: 10μm. **(B-D)** The area occupied by ubiquitin and ubiquitin intensity were measured and plotted over time. Cells were analyzed at 0h (untreated), 3h puromycin, and 3h-post washout with either media (B), media + CQ (C), or media + MG132 (D). Each symbol represents the mean of n=4 experiments ± SD (23-40 cells per experiment). Significant *p* values comparing cell lines are noted (unpaired *t* tests).

To investigate how WHAMM/JMY^DKO^ cells might be degrading ubiquitin foci via the ALS and UPS, we performed the puromycin treatment and washout experiments using recovery media containing either chloroquine to inhibit lysosome-mediated degradation, or MG132 to inhibit proteasome-mediated degradation. Upon recovery in media containing chloroquine, the ubiquitin staining in eHAP cells increased (Figure 3A). The ubiquitin structures that remained were smaller than the foci that were present before drug washout, but they collectively occupied a larger area of the cell and exhibited a greater intensity (Figure 3C). In contrast, recovery in chloroquine had only a modest effect on ubiquitin staining in WHAMM/JMY^DKO^ cells (Figure 3A). The cell area occupied by ubiquitin was unchanged, while the intensity of the structures slightly decreased (Figure 3C). Taken together with the reduced LC3-II levels observed in DKO cells, these results suggest that WHAMM/JMY^DKO^ cells do not utilize the ALS as their primary means for degrading ubiquitinated cargo.

We thus turned our attention to the UPS. Upon recovery in media containing MG132, the ubiquitin staining occupied a significantly larger area in WHAMM/JMY^DKO^ cells than in eHAP cells, and the staining was significantly more intense (Figure 3, A and D). Moreover, the DKO cells sometimes contained an extremely bright singular aggresome-like ubiquitin structure adjacent to the nucleus (Figure 3, A and D). Overall, these data suggest that wild type cells can effectively utilize the ALS to degrade ubiquitinated cargo during proteotoxic stress, whereas WHAMM/JMY-deficient cells are more reliant on the UPS to maintain proteostasis.

We hypothesized that proteasomes would be active at ubiquitin foci, so we tested whether these structures were sites of proteasome-mediated degradation by treating the eHAP and DKO cells with fresh media or media containing puromycin for 3h and adding the proteasome activity probe Me4BodipyFL-Ahx3Leu3VS prior to fixation (Verdoes *et al*., 2006). In the absence of puromycin, the probe was diffuse and dim in both cell lines (Figure 4A). However, after puromycin exposure, the probe became enriched in the ubiquitin foci, which were much more prevalent in WHAMM/JMY^DKO^ cells (Figure 4A). These results further support the idea that the WHAMM/JMY^DKO^ cells depend on the UPS for misfolded protein degradation.

**Figure 4.**
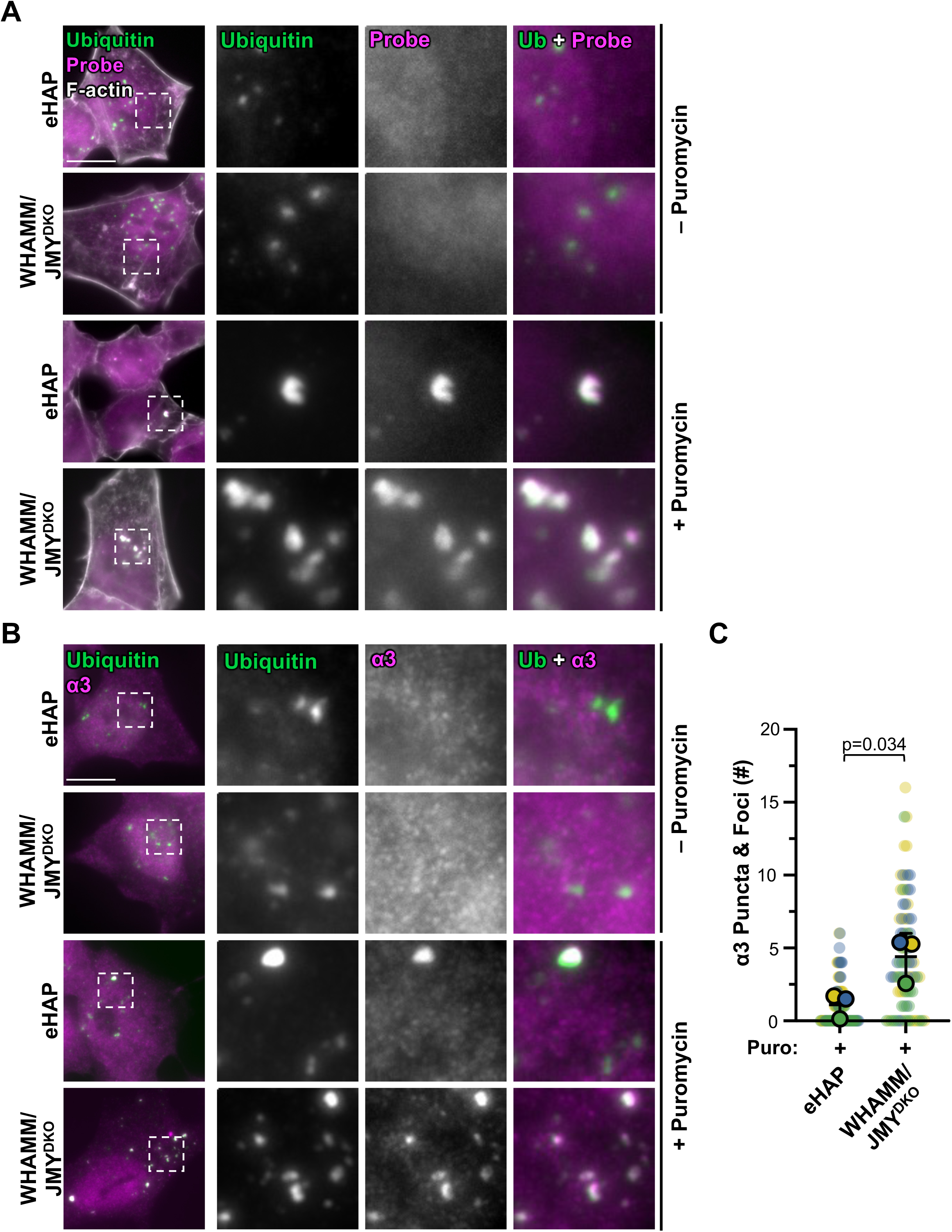
WHAMM/JMY^DKO^ cells have more active proteasomes at ubiquitin foci and puncta. **(A)** eHAP and WHAMM/JMY^DKO^ cells were treated with 2μg/mL puromycin for 3h, with the proteasome activity probe (magenta) added 30min before being fixed and stained with an antibody to visualize ubiquitin conjugated proteins (green) and phalloidin to visualize F-actin (white). Scale bar: 10μm. **(B)** eHAP and WHAMM/JMY^DKO^ cells were treated with 2μg/mL puromycin for 3h and stained with antibodies to visualize ubiquitin-conjugated proteins (green) and the α3 subunit of the proteasome (magenta). Scale bar: 10μm. **(C)** The number of α3 puncta and foci were counted per cell. Each small symbol represents a single cell (26-32 cells per experiment), each larger symbol represents the experimental average from n=3 replicates ± SD. Each experiment is coordinated by color. Significant *p* value is noted (unpaired *t* test).

To examine the localization of proteasomes in eHAP and WHAMM/JMY^DKO^ cells, we stained them with antibodies to ubiquitin and to the α3 or α6 subunits of the 20S core proteasomal particle. At steady state, α3 and α6 staining was diffuse throughout the nucleus and cytosol in both cell lines, with no noticeable enrichment at small ubiquitin puncta (Figure 4B and Supplemental Figure S3). After puromycin treatment, α3 and α6 staining was abundant at ubiquitin puncta as well as the larger, brighter ubiquitin foci (Figure 4B and Supplemental Figure S3). Quantification of the number of α3 puncta and foci demonstrated that WHAMM/JMY^DKO^ cells had approximately 4-fold more proteasome-associated structures compared to eHAP cells (Figure 4C). In contrast, staining with antibodies to LC3 did not reveal any enrichment at ubiquitin foci in either cell line, although LC3 was present at some smaller ubiquitin structures in eHAP cells (Supplemental Figure S4). Together, the activity and immunolocalization results indicate that the WHAMM/JMY-deficient cells redistribute proteasomes but do not assemble autophagosomes in order to counteract proteotoxic stress.

### Synphilin-1 is enriched in WHAMM/JMY^DKO^ cells and localizes with ubiquitinated proteins

Given the LC3 lipidation deficiency in WHAMM/JMY^DKO^ cells and their reliance on the UPS for removing misfolded polypeptides, we wondered whether stochastic fluctuations in stress that occur under normal culturing conditions might have led to adaptive, proteostasis-maintaining changes in the DKO cell transcriptome. To address this question, we performed RNA-sequencing (RNA-seq) and differential gene expression analyses on parental eHAP and WHAMM/JMY^DKO^ cells. Since the DKO cell line was derived by first creating a WHAMM knockout cell line and then inactivating JMY (King *et al*., 2021), we additionally included WHAMM^KO^ cells in these analyses.

Upon filtering for genes whose expression differed from that in eHAP cells by at least 1.5-fold and that were statistically significant (q<0.05), we found that the abundance of approximately 0.6% of mapped transcripts were affected by the deletion of WHAMM and 1.3% by the WHAMM/JMY double deletion (Figure 5A). Among these, significant changes were not observed in other actin nucleation factors, autophagy-related genes such as autophagy receptors or lysosomal components, or UPS genes such as proteasome subunits or chaperones (Supplemental Data File). Since WHAMM and JMY both function in autophagy, we were most interested in identifying genes that might be up-regulated in a stepwise fashion in the single and double knockout cells to potentially compensate for impaired ALS-mediated proteostasis. Only six genes fit this category: *ZNF300*, *POSTN*, *COL5A2*, *FAM84B*, *TPM2*, and *SNCAIP* (Figure 5A). Among these, we focused on *SNCAIP*, which has previously been shown to impact proteostasis in the context of Parkinson’s Disease.

**Figure 5.**
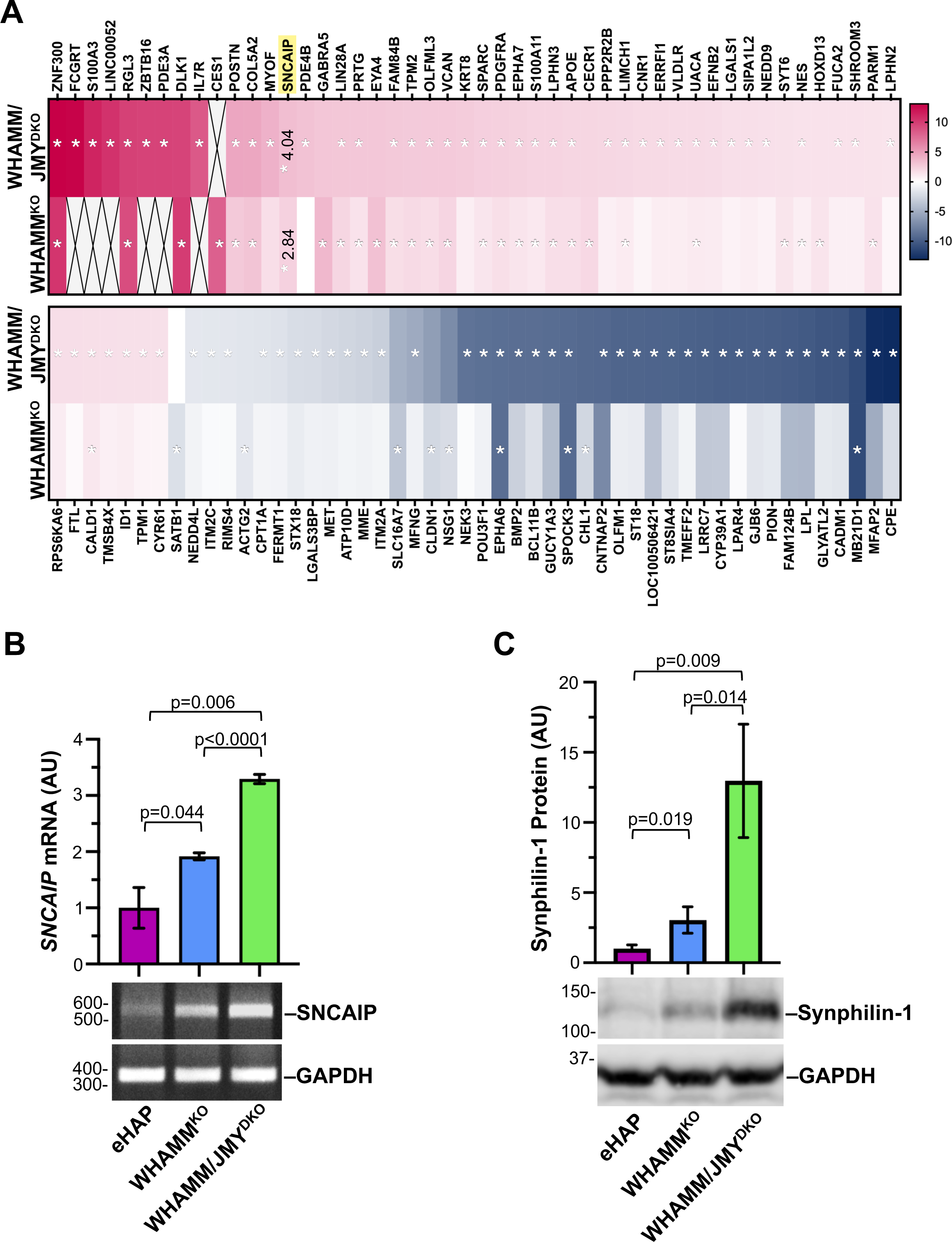
WHAMM and WHAMM/JMY deletions change cellular gene expression resulting in a stepwise upregulation of the *SNCAIP* gene and an increased abundance of the Synphilin-1 protein. **(A)** RNA collected from eHAP, WHAMM^KO^, and WHAMM/JMY^DKO^ cells was subjected to mRNA sequencing analyses. The heatmap reflects fold change in the fragments per kilobase mapped (FPKM) in KO vs. parental cells, expressed as log_2_(KO FPKM : eHAP FPKM), of genes that are at least 1.5-fold different from the parental eHAP. Asterisks denote values that are statistically significant (*q*<0.05) in one or more of the cell lines. Boxes marked with X denote genes with undetected transcripts in their respective dataset. Log_2_(KO FPKM : eHAP FPKM) values for *SNCAIP* are noted. **(B)** RNA collected from eHAP, WHAMM^KO^, and WHAMM/JMY^DKO^ cells was subjected to RT-PCR with primers to *SNCAIP* and *GAPDH*. *SNCAIP* band intensities were normalized to *GAPDH* and quantified, and each bar represents the average from n=3 experiments ± SD. Significant *p* values are noted (unpaired *t* tests). **(C)** eHAP, WHAMM^KO^, and WHAMM/JMY^DKO^ cells were subjected to SDS-PAGE and immunoblotting with antibodies to Synphilin-1 and GAPDH. Synphilin-1 band intensities were normalized to GAPDH and quantified, and each bar represents the average from n=3 experiments ± SD.

*SNCAIP* encodes the protein Synphilin-1, which was originally discovered as an α-synuclein binding partner (Engelender *et al*., 1999), and soon after found to be present in Lewy Bodies (Wakabayashi *et al*., 2000). Synphilin-1 has been proposed to play multiple roles in proteostasis pathways. It may interact with the regulatory 19s particle of the proteasome (Marx *et al*., 2007), cooperate with α-synuclein to recruit LC3 and LAMP1 for autophagic degradation of protein aggregates (Wong *et al*., 2008), and form aggresome-like inclusions (Zaarur *et al*., 2008; Wong *et al*., 2012). To corroborate the RNA-sequencing results, we performed RT-PCR with primers to *SNCAIP* and confirmed stepwise increases in mRNA levels in WHAMM^KO^ and WHAMM/JMY^DKO^ cells (Figure 5B). *SNCAIP* was not up-regulated when JMY was knocked out by itself (King *et al*., 2021).

To determine if the mRNA levels in eHAP, WHAMM^KO^, and WHAMM/JMY^DKO^ cells were translated into differences in protein abundance, we prepared whole cell extracts and immunoblotted the samples using antibodies to Synphilin-1. Compared to parental eHAP cells, WHAMM^KO^ cells expressed approximately 3-fold more Synphilin-1 (Figure 5C). In the WHAMM/JMY^DKO^ cells, Synphilin-1 levels were quite variable, but averaged about 4-fold more than what was found in the single WHAMM^KO^ cells, and up to 12-fold more than the amount in eHAP cells (Figure 5C). These results show that upon sequential loss of WHAMM and JMY, cells produce more of both the *SNCAIP* transcript and the Synphilin-1 protein.

Because of the large increase in Synphilin-1 abundance in WHAMM/JMY^DKO^ cells compared to the original eHAP parental cells, we used these two cell lines to probe the localization of Synphilin-1 and ubiquitin both at steady state and in the presence of puromycin. Following fixation and staining with antibodies to ubiquitin and Synphilin-1, the two proteins were found in overlapping punctate structures in both eHAP and WHAMM/JMY^DKO^ cells (Figure 6A). Clusters of Synphilin-1 puncta often localized around the edges of ubiquitin puncta, rather than colocalizing with the centers of the ubiquitin structures themselves (Figure 6A). Approximately 50-60% of Synphilin-1 staining overlapped with ubiquitin staining in the absence or presence of puromycin (Figure 6B). Fluorescent intensity plot profiles showed that Synphilin-1 puncta formed around the smaller, dimmer ubiquitin puncta, but not around bright ubiquitin foci (Figure 6, C and D). These results indicate that endogenous Synphilin-1 associates with ubiquitin-conjugated protein aggregates, but not densely ubiquitinated foci, and that this relationship takes place whether or not cells express WHAMM and JMY.

**Figure 6.**
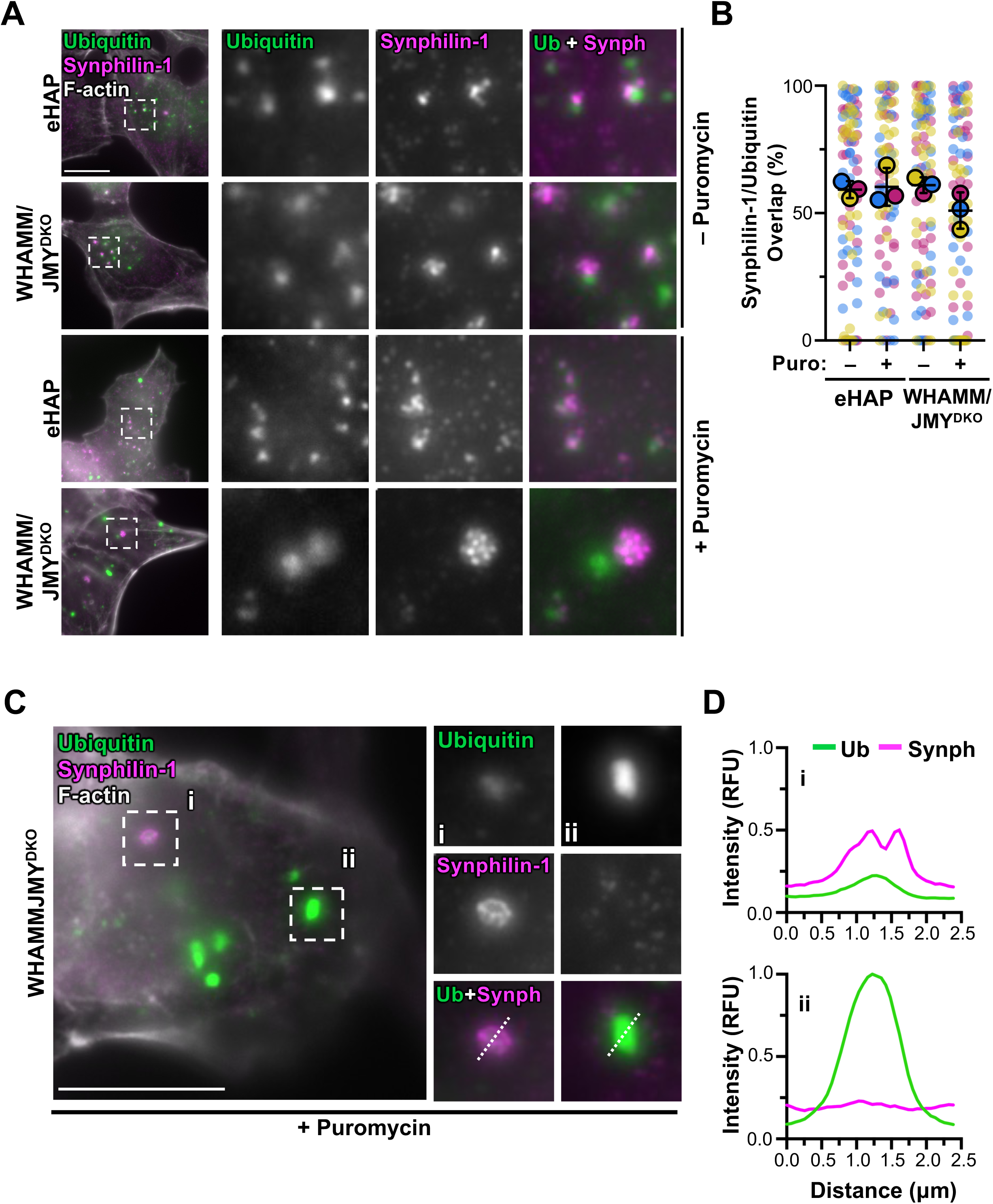
Synphilin-1 clusters associate with ubiquitin puncta. **(A)** eHAP and WHAMM/JMY^DKO^ cells were treated with 2μg/mL puromycin for 3h and stained with antibodies to visualize ubiquitin-conjugated proteins (green), Synphilin-1 (magenta), and with phalloidin to visualize F-actin (white). Magnifications represent examples of Synphilin-1 clusters around ubiquitin puncta. Scale bar: 10μm. **(B)** The percentage of Synphilin-1 staining overlapping with ubiquitin staining in puromycin treated cells was calculated and Mander’s coefficients were generated. Each small point represents one cell (29-32 cells per experiment), each larger symbol represents the experimental average from n=3 replicates ± SD. Each experiment is coordinated by color. **(C)** WHAMM/JMY^DKO^ cells were treated with 2μg/mL puromycin for 3h and stained with antibodies to visualize ubiquitin-conjugated proteins (green), Synphilin-1 (magenta), and with phalloidin to visualize F-actin (white). Scale bar: 10μm. Magnifications show the clustering of Synphilin-1 (i) at dimmer ubiquitin puncta and (ii) not at larger ubiquitin foci. **(D)** 2.5μm lines were drawn through ubiquitin structures in (C) to measure pixel intensity profiles. Ubiquitin intensities were normalized to inset (ii), and Synphilin-1 intensities were normalized to inset (i).

### Synphilin-1 overexpression can drive condensate formation via its ANK1 domain

To explore the association between Synphilin-1 and ubiquitinated proteins in a tractable live cell system, we generated Cos7 cells stably expressing mCherry-Synphilin-1 (Supplemental Figure S5) and transiently transfected them with a plasmid encoding GFP-Ubiquitin prior to timelapse imaging. Even without puromycin treatment, cells expressing GFP-Ubiquitin readily formed cytosolic puncta and foci (Figure 7A), consistent with previous observations of ubiquitin overexpression (Dantuma *et al*., 2006; Saini *et al*., 2021). Live imaging demonstrated that both mCherry-Synphilin-1 and GFP-Ubiquitin structures consistently moved throughout the cytosol in a dynamic fashion (Figure 7A and Supplemental Video S1). mCherry-Synphilin-1 and GFP-Ubiquitin puncta were frequently in close proximity to one another, and often remained paired during movement even when GFP-Ubiquitin appeared to change in shape. mCherry-Synphilin-1 and GFP-Ubiquitin also formed larger structures that changed morphology before separating into two pieces (Figure 7A, Supplemental Video S1). These live studies reveal that Synphilin-1 is capable of associating with and moving concurrently with ubiquitinated cargo.

**Figure 7.**
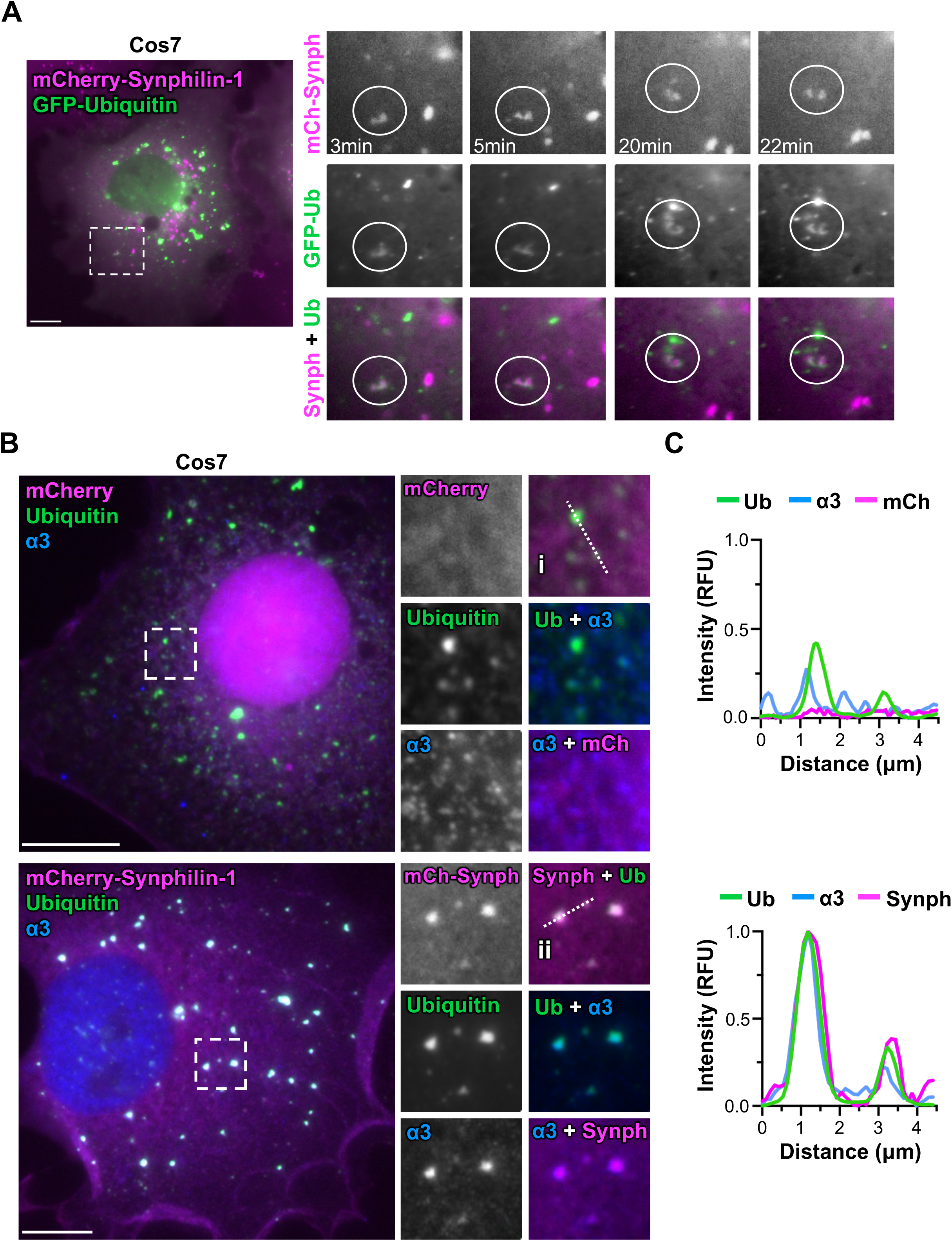
Synphilin-1 co-migrates with ubiquitinated cargo undergoing morphological changes and localizes together with ubiquitin and proteasomes. **(A)** Cos7 cells stably expressing mCherry-Synphilin-1 (magenta) were transfected with plasmids encoding GFP-Ubiquitin (green) before live imaging. Scale bar: 10μm. **(B)** Cos7 cells stably expressing mCherry or mCherry-Synphilin-1 (magenta) were treated with 5μg/mL puromycin for 2h and stained with antibodies to ubiquitinated conjugates (green) and the α3 subunit of the proteasome (blue). Magnifications show ubiquitin structures in control cells or highlight structures exhibiting mCherry-Synphilin-1, ubiquitin, and α3 colocalization. Scale bars: 10μm. **(C)** 4.5μm lines were drawn through structures in (B) to generate pixel intensity profiles. Intensities from (i) were normalized to each corresponding channel in (ii).

We next wanted to dissect the relationship between the domains of Synphilin-1 and its localization to endogenous ubiquitinated proteins. We cloned GFP-fusion constructs containing full-length Synphilin-1 and a series of truncations that include the N-terminal region (NT); the N-terminus plus first Ankyrin repeat and central coiled-coil domain (NT-ANK1-CC); the coiled-coil, second Ankyrin repeat, and C-terminus (CC-ANK2-CT); and the C-terminal region (CT) (Figure S5). Similar Synphilin-1 constructs were previously shown to form aggresome-like inclusions and aggregates in mouse and human cells depending on the presence of either Ankyrin repeat and the CC domain (Zaarur *et al*., 2008; Wong *et al*., 2012). We transiently expressed the fusions in Cos7 cells, fixed the cells, and observed the fluorescent localization patterns (Figure S5). Full length Synphilin-1 formed small puncta, bright foci, and large inclusions, highlighting the 3 types of localization that we previously observed with the endogenous protein in WHAMM/JMY^DKO^ cells. Among the truncations, the NT fragment was diffuse throughout the cytosol while the CT was present in the cytosol and nucleus. As previously shown (Wong *et al*., 2012), the ANK1-CC construct accumulated in large, intense cytosolic inclusions whereas the CC-ANK2 construct localized in smaller puncta.

We also monitored the dynamics of full-length GFP-Synphilin-1 by live imaging. As observed in cells stably expressing mCherry-Synphilin-1, GFP-Syphilin-1 structures were dynamic and moved throughout the cytosol. Sometimes Synphilin-1 formed bright foci that coalesced to form juxtanuclear inclusions (Supplemental Video S2), while other times foci broke down over time (Supplemental Video S3). Because of the intensity and size of the inclusions formed by full length Synphilin-1 and the ANK1-CC domains, we wanted to determine if these structures had liquid-like properties, so we treated the GFP-Synphilin-1 transfected Cos7 cells with either 0.3M NaCl or 0.4M Sorbitol as a means for disrupting cytosolic condensates prior to fixation. Instead of the large, round inclusions that formed in untreated cells, smaller and more numerous cytosolic structures were distributed throughout the osmotically manipulated cells (Supplemental Figure S5). To confirm the localization of the Ankyrin repeats to ubiquitinated proteins, we treated GFP-NT-ANK1-CC- or GFP-CC-ANK2-CT-expressing cells with puromycin before staining them with ubiquitin antibodies. Most of the cytosolic NT-ANK1-CC foci were enriched with ubiquitin, as were some of the CC-ANK2-CT puncta (Supplemental Figure S5). These results are consistent with previous suggestions that Synphilin-1 primarily utilizes its first Ankyrin repeat and CC domain for the formation of ubiquitin-associated condensates.

### Synphilin-1 coordinates proteasome-mediated removal of ubiquitinated proteins and p62

The prominence of Synphilin-1, ubiquitinated proteins, and proteasomes in both genetically-modified Cos7 cells and intrinsically-altered WHAMM/JMY^DKO^ cells led us to next explore the simultaneous localization of these factors as well as the impact of Synphilin-1 on proteostasis. To see which Synphilin-1 and ubiquitin structures were accompanied by proteasomes, we first examined α3 proteasome subunit immunolocalization in puromycin-treated mCherry-Synphilin-1-expressing Cos7 cells. In this exogenous expression system, all three proteins localized to the same structures (Figure 7B), and fluorescence intensity profiles highlighted their presence in both bright cytosolic foci and dimmer puncta (Figure 7, B and C). In contrast, cells expressing mCherry formed some ubiquitin foci and puncta, but mCherry was not enriched at these structures (Figure 7, B and C).

The tripartite localization of ubiquitin and proteasomes with Synphilin-1 was a reminder that SQSTM1/p62 is another protein which localizes to cytoplasmic bodies and functions in proteostasis. p62 acts as an autophagy receptor that links ubiquitinated cargo to LC3 in autophagosomes, interacts with proteasomes to influence UPS activity, and forms protein condensates (Seibenhener *et al*., 2004; Myeku and Figueiredo-Pereira, 2011; Cohen-Kaplan *et al*., 2016; Danieli and Martens, 2018; Sun *et al*., 2018; Zaffagnini *et al*., 2018; Fu *et al*., 2021; 2024; Feng *et al*., 2022). This led us to question whether p62 was found in the same cytosolic structures as ubiquitin and Synphilin-1. In GFP-Synphilin-1-expressing Cos7 cells, anti-ubiquitin and anti-p62 immunostaining revealed that GFP-Synphilin-1 foci and puncta frequently overlapped with both ubiquitin and p62 at steady state and upon treatment with puromycin (Figure 8A). As with mCherry earlier, GFP by itself did not coalesce into foci or puncta in puromycin-treated cells as GFP-Synphilin-1 and p62 did (Figure 8A), indicating that not all proteins are incorporated into cytoplasmic bodies in puromycin-treated cells.

**Figure 8.**
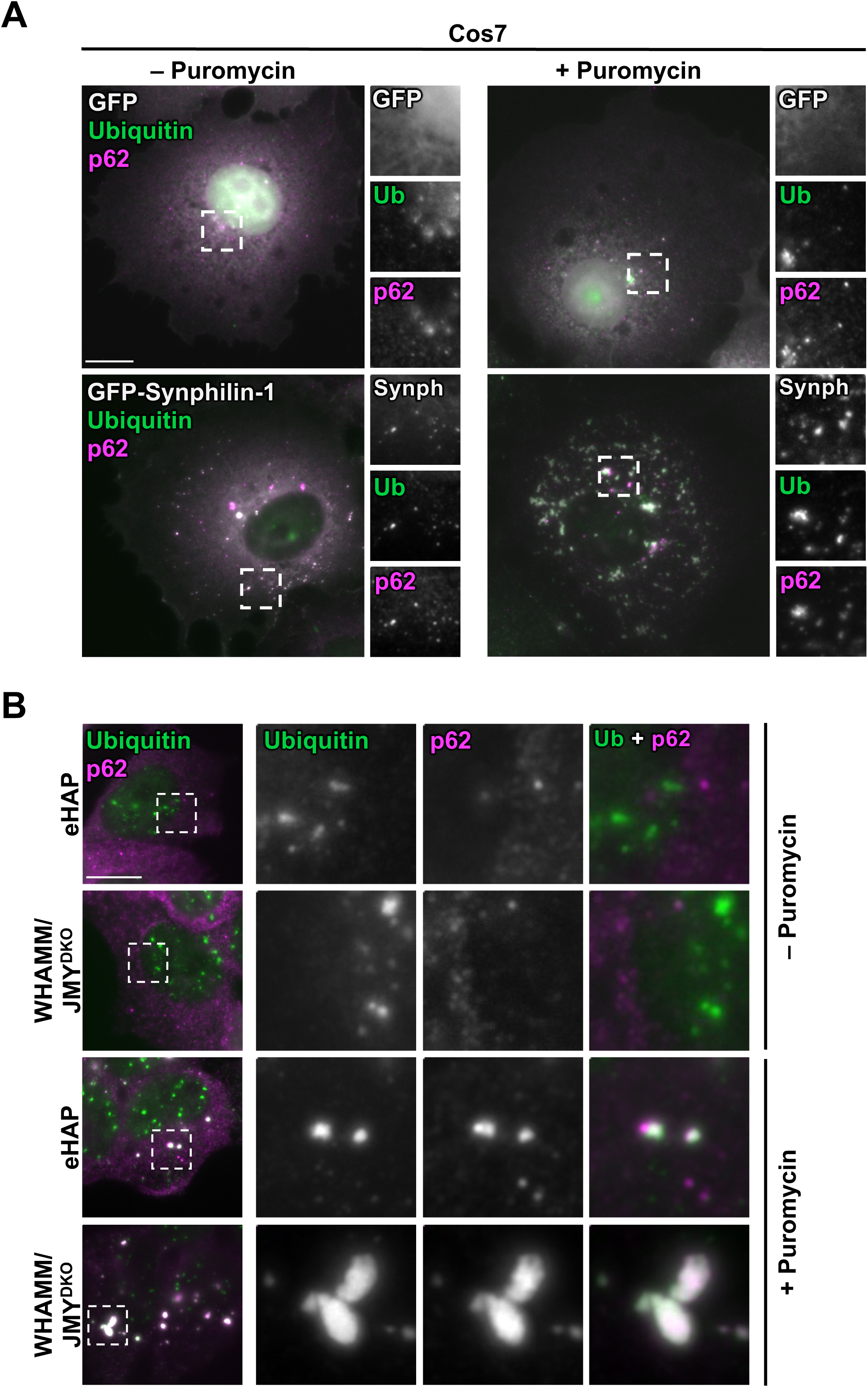
SQSTM1/p62 localizes to ubiquitin foci and GFP-Synphilin-1 structures. **(A)** Cos7 cells were transfected with GFP or GFP-Synphilin-1 (white), treated with 2μg/mL puromycin for 3h, and stained with antibodies to ubiquitinated proteins (green) and p62 (magenta). Magnifications represent examples of ubiquitin-p62 overlap. Scale bar: 10μm. **(B)** eHAP and WHAMM/JMY^DKO^ cells were treated with 2μg/mL puromycin for 3h and stained with antibodies to visualize ubiquitinated proteins (green) and p62 (magenta). Scale bar: 10μm.

To define the localization of p62 in relation to ubiquitin in the endogenous Synphilin-1 up-regulation system, we also examined eHAP and WHAMM/JMY^DKO^ cells using immunofluorescence. Under normal culture conditions, endogenous nuclear ubiquitin puncta did not colocalize with p62 in either cell line (Figure 8B). However, after cells were exposed to puromycin, ubiquitinated proteins and p62 overlapped within cytoplasmic foci, including the larger, brighter structures that formed in the DKO cells (Figure 8B).

Finally, we sought to determine the functional consequences of Synphilin-1 upregulation in WHAMM/JMY^DKO^ cells by targeting Synphilin-1 for depletion using two independent siRNAs (siSYN^A^, siSYN^B^) to the *SNCAIP* transcript. In eHAP cells, the siRNAs reduced the already-low amounts of Synphilin-1 to barely detectable levels, and in WHAMM/JMY^DKO^ cells were able to decrease the high Synphilin-1 levels by ∼75% (Supplemental Figure S6).

Based on our earlier experiments assessing cell death and ubiquitinated protein accumulation at different times, we chose 7h as a single intermediate timepoint to effectively capture the cell viability and proteostasis phenotypes of Synphilin-1-depleted cells. The combination of Synphilin-1 depletion and puromycin exposure increased the frequency of PI positivity and apoptotic body formation in both the parental and WHAMM/JMY-deficient cells, with the Synphilin-1-depleted WHAMM/JMY^DKO^ samples exhibiting the most death (Supplemental Figure S6). We also noticed that the DKO cells, especially those without Synphilin-1, displayed a higher frequency of micronuclei (Supplemental Figure S6), cytoplasmic chromatin structures that can arise from mitotic errors and DNA damage (Zhao *et al*., 2023). These results are consistent with the conclusion that Synphilin-1 up-regulation in WHAMM/JMY^DKO^ cells plays a cytoprotective role.

To determine the effects of Synphilin-1 on proteostasis, we exposed control or Synphilin-1-depleted eHAP and WHAMM/JMY^DKO^ cells to puromycin, and then immunoblotted the cell extracts and quantified the band intensities of Synphilin-1, p62, and high molecular weight ubiquitinated proteins relative to tubulin and GAPDH loading controls (Figure 9A-C). While the two different *SNCAIP* siRNAs gave different degrees of Synphilin-1 knockdown (Figure 9 and Supplemental Figure S7), several patterns of p62 and ubiquitinated protein abundance emerged. Relative to samples treated with control siRNAs, the more effective *SNCAIP* siRNA (siSYN^B^) caused a 50% increase in p62 levels in eHAP cells and 200% increase in p62 levels in DKO cells (Figure 9, A and B). Plotting the p62 densitometry values against the Synphilin-1 knockdown percentages from the different siRNAs across multiple experiments revealed a positive correlation between p62 accumulation and Synphilin-1 depletion, with the WHAMM/JMY^DKO^ data showing a stronger relationship (Figure 9D). Similar trends in p62 levels were observed in cells that were not exposed to puromycin, although the magnitudes of p62 increase were lower (Supplemental Figure S7). These observations further imply that WHAMM/JMY-deficient cells are more sensitive than eHAP cells to Synphilin-1 depletion and accumulate the p62 adaptor protein, particularly during puromycin-induced proteotoxic stress.

**Figure 9.**
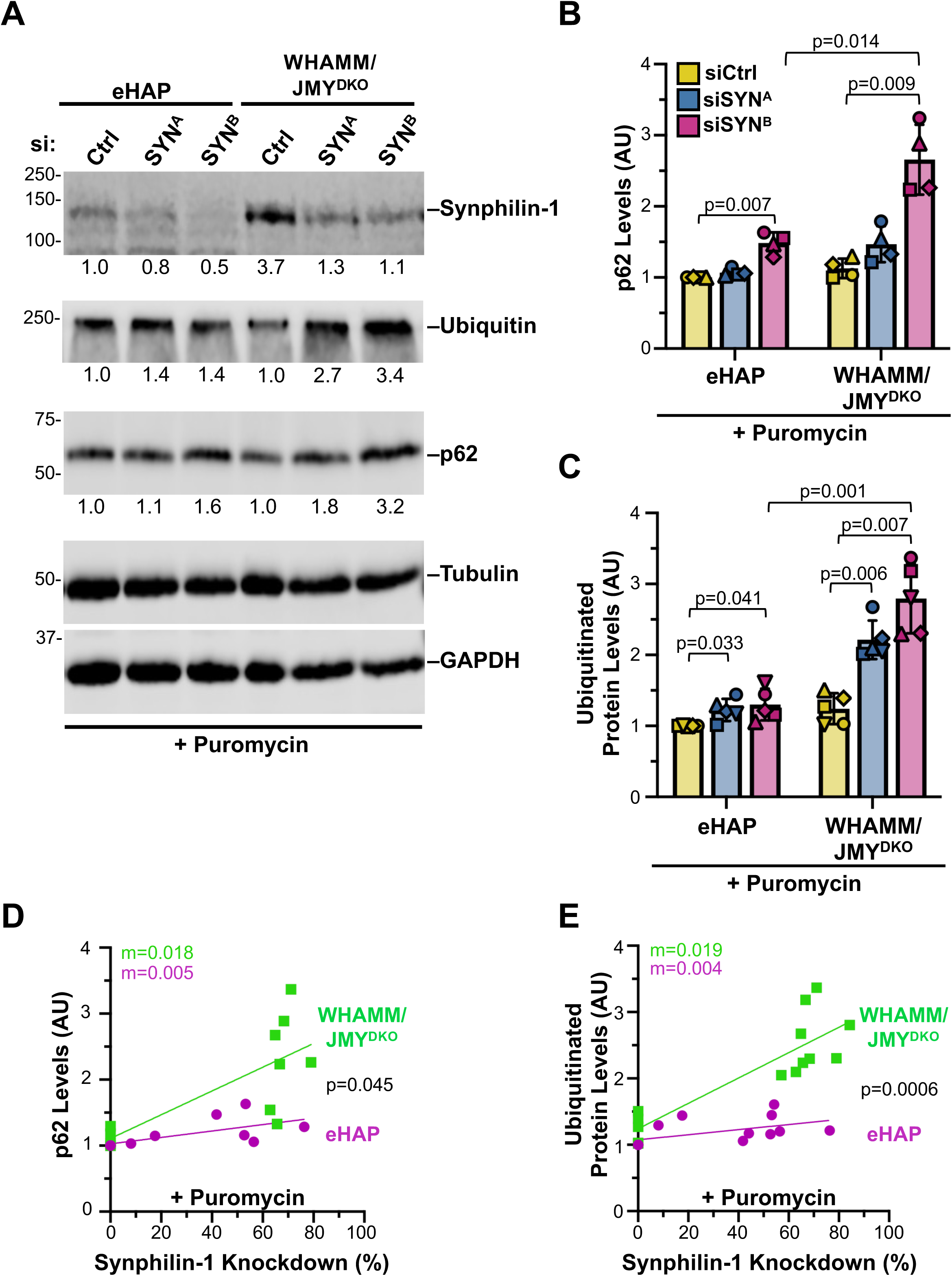
Synphilin-1 limits the accumulation of ubiquitinated proteins in WHAMM/JMY^DKO^ cells. **(A)** eHAP and WHAMM/JMY^DKO^ cells were transfected with control siRNAs (siCtrl) or siRNAs targeting the *SNCAIP* transcript (siSYN), then treated with 1μg/mL puromycin for 7h before immunoblotting with antibodies to Synphilin-1, ubiquitinated proteins, p62, Tubulin, and GAPDH. Densitometry values for Synphilin-1, high molecular weight ubiquitin, and p62 bands were normalized to Tubulin and GAPDH controls and are shown beneath their respective blots. **(B)** p62 band densitometry values were normalized to Tubulin and GAPDH loading controls and quantified. Each bar represents the mean from n=4 experiments ± SD. Significant p-values are noted (paired *t* tests for comparisons within the same cell line, and unpaired *t* tests for comparisons between cell lines). **(C)** Ubiquitin band densitometry values were normalized to Tubulin and GAPDH loading controls and quantified. Each bar represents the mean from n=5 experiments ± SD. Significant p-values are noted (paired *t* tests for comparisons within the same cell line, and unpaired *t* tests for comparisons between cell lines). **(D)** Densitometry values of p62 were plotted against the percentage of Synphilin-1 knockdown (calculated by dividing the densitometry value of Synphilin-1 in siSYN cells by the value of Synphilin-1 in siCtrl cells). Each point represents one sample, with samples from both siSYN^A^ and siSYN^B^ treatments shown. The slopes (m) in the linear trendline regression equations for eHAP (Y = 0.004865x + 1.027) and WHAMM/JMY^DKO^ (Y = 0.01801x + 1.108) were significantly nonzero (p=0.031 and p=0.0073 respectively). The 3.7-fold difference between the slopes was statistically significant (p=0.045). **(E)** Densitometry values of ubiquitin were plotted against the percentage of Synphilin-1 knockdown. Each point represents one sample, with samples from both siSYN^A^ and siSYN^B^ shown. The slopes (m) in the linear trendline regression equations for eHAP (Y = 0.003838x + 1.075) and WHAMM/JMY^DKO^ (Y = 0.01911x + 1.242) were significantly nonzero (p=0.0493 and p<0.0001 respectively). The 5-fold difference between the slopes was statistically significant (p=0.0006).

When protein ubiquitination levels were evaluated, both independent *SNCAIP* siRNAs caused significant phenotypes. In eHAP cells, the amounts of high molecular weight ubiquitin conjugates were elevated by about 25% relative to control siRNA-treated cells, while in WHAMM/JMY^DKO^ samples the ubiquitination levels were 2-3-fold higher, depending on the potency of the siRNA (Figure 9, A and C and Supplemental Figure S7). Comparisons of ubiquitinated protein levels to Synphilin-1 knockdown percentages indicated that WHAMM/JMY^DKO^ cells had a stronger positive correlation than eHAP cells (Figure 9E). Thus, WHAMM/JMY-deficient cells require high levels of Synphilin-1 in order to efficiently manage ubiquitinated protein removal during proteotoxic stress.

## DISCUSSION

Proteostasis is critical for cellular and organismal health, but how cells cope with proteotoxic stress when their autophagosome-lysosome system is compromised is not well understood. Consistent with previous work showing that the Arp2/3 complex and its activators WHAMM and JMY function at autophagosomes and lysosomes, our current study establishes that WHAMM and JMY double knockout cells have a defect in LC3 lipidation and autophagic degradation. We further demonstrate that these cells have adapted to rely on the ubiquitin-proteasome system as a means of mitigating the accumulation of truncated proteins. Moreover, up-regulation of Synphilin-1 allowed the cells to better modulate protein aggregation, promote proteasome-mediated degradation, and protect against cytotoxicity (Figure 10). These results highlight the versatile mechanisms that allow cells to maintain proteostasis.

**Figure 10.**
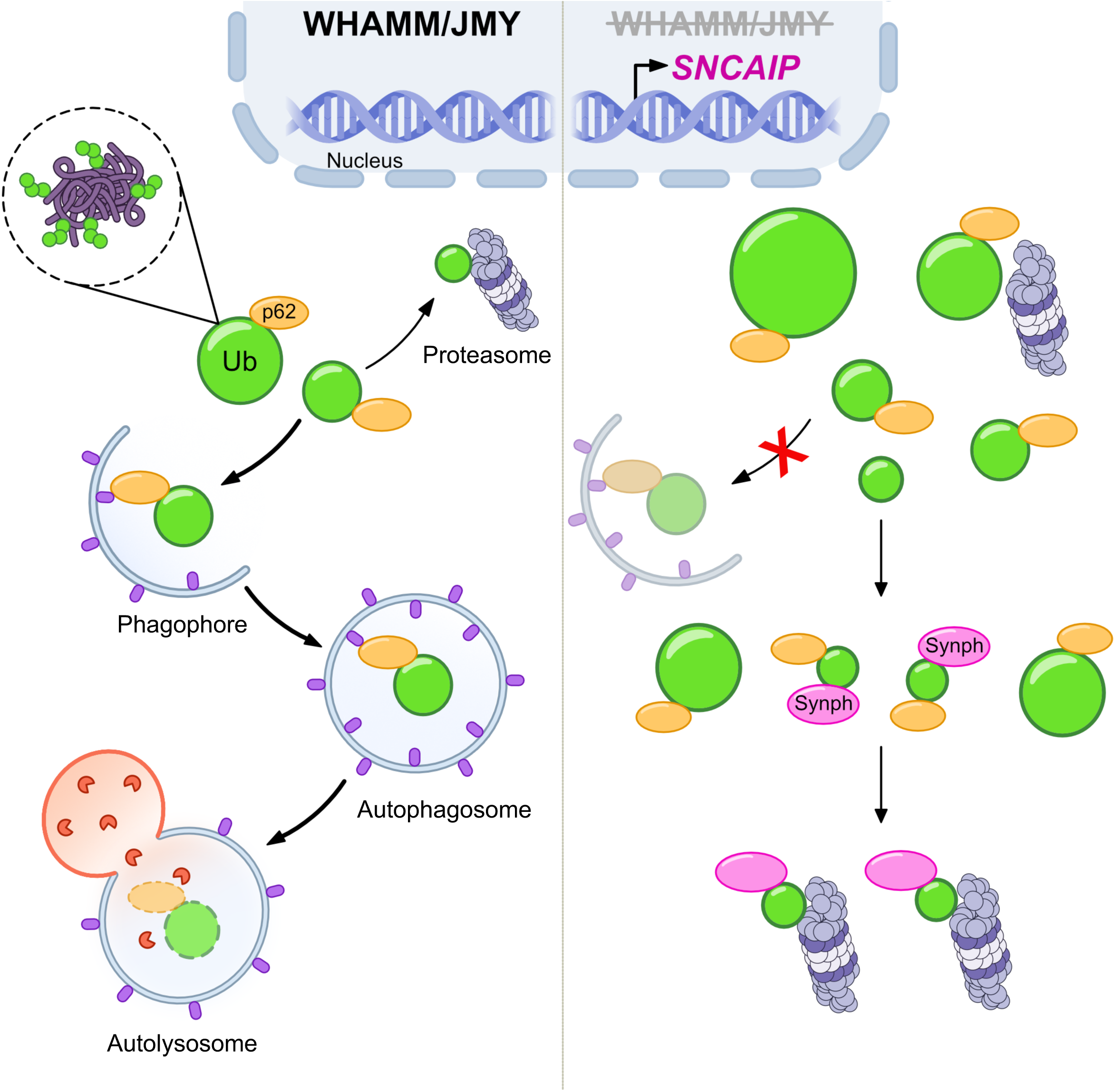
Synphilin-1 and the Ubiquitin Proteasome System cooperate to maintain proteostasis during stress. Left: Cells with WHAMM, JMY, and a functional autophagy pathway form foci and puncta (green circles) composed of ubiquitinated misfolded proteins, during proteotoxic stress. Substrates can be targeted for proteasomal degradation or to autophagosomes via autophagy receptors such as SQSTM1/p62 for lysosomal degradation. Right: WHAMM/JMY- and autophagy-deficient cells accumulate large, intense foci that can be engaged by p62, but are unable to be targeted to the Autophagosome Lysosome System. These cells increase expression of Synphilin-1, which localizes with p62, proteasomes, and ubiquitinated cargo to promote their proteasome-mediated degradation, maintain proteostasis, and enable cell survival. Model made using Affinity Designer. DNA icons retrieved from NIAID NIH BioArt Source (bioart.niaid.nih.gov/bioart/123).

The influence of the actin cytoskeleton on the proteome begins at chromatin remodeling and gene transcription, and ends with protein degradation via the ALS or UPS (Williams and Rousseau, 2022), and the specific contributions of actin regulators to proteostasis have started to come into focus in recent years. Many actin nucleation factors, including WHAMM and JMY, along with WASP, WASH, and Cortactin, function in various steps of the ALS in diverse cellular contexts (Campellone *et al*., 2023). However, the role of the cytoskeleton in the UPS is less well understood. Actin has been shown to associate with proteasomes in yeast and human fibroblasts (Arcangeletti *et al*., 1997; Haarer *et al*., 2011), and it regulates the assembly of proteasome subunits in yeast (Williams *et al*., 2022). The link between actin and the proteasome itself or its clients is an open avenue for investigation. It would be interesting to determine if the Arp2/3 complex or WASP-family members generate actin filaments that associate with the proteasome.

WHAMM and JMY constitute a subgroup of nucleation-promoting factors in the WASP-family and function at distinct steps in multiple fundamental cellular processes ranging from anterograde transport (Campellone *et al*., 2008; Schlüter *et al*., 2014; Russo *et al*., 2016), to autophagy (Coutts and La Thangue, 2015; Kast *et al*., 2015; Mathiowetz *et al*., 2017), and to apoptosis (King *et al*., 2021; King and Campellone, 2023). WHAMM/JMY^DKO^ cells are viable and do not have any obvious growth abnormalities under standard culturing conditions, although exposure to genotoxic stress previously revealed that WHAMM/JMY-deficient cells are defective at intrinsic apoptotic signaling (King *et al*., 2021; King and Campellone, 2023). In the current study, we found that the DKO cells are more sensitive to proteotoxic stress and accumulate ubiquitinated proteins in cytoplasmic foci after exposure to puromycin. Surprisingly, cells lacking WHAMM and JMY are still able to eliminate the ubiquitin foci after removal of puromycin, despite their deficiency in LC3-I to LC3-II conversion. Our washout studies using lysosome or proteasome inhibitors, coupled with our assessments of proteasomal localization and activity at ubiquitin foci and puncta, revealed a unique dependency on the UPS in this cellular context.

While WHAMM/JMY^DKO^ cells did not appear to target ubiquitinated protein aggregates or condensate-like foci to autophagosomes, they are presumably able to employ disaggregation factors, chaperones, and shuttling factors to extract proteins and target them to the proteasome for degradation (Hjerpe *et al*., 2016; Mauthe *et al*., 2025).

To ascertain whether compensatory gene expression changes might play a role in WHAMM/JMY^DKO^ cell survival and response to stress, we performed mRNA sequencing on parental eHAP, WHAMM^KO^, and WHAMM/JMY^DKO^ cells. By filtering for genes that underwent a stepwise up-regulation from the parental cell line to a single WHAMM KO to the compound JMY KO, we sought to find candidates that might help cells adapt to the sequential loss of WHAMM and JMY. While five of the six genes that fit this expression pattern – *ZNF300*, *POSTN*, *COL5A2*, *FAM84B*, and *TPM2* – may provide interesting links to transcriptional regulation, extracellular matrix remodeling, and actin filament functionality (Qiu *et al*., 2008; Liu *et al*., 2014; Gunning *et al*., 2015; Walker *et al*., 2018; Huang *et al*., 2021), we were drawn to the presence of *SNCAIP*, which encodes the protein Synphilin-1.

Synphilin-1 was discovered in a screen for α-synuclein-interacting proteins, and later found to be present with α-synuclein in Lewy Bodies and at synaptic terminals (Engelender *et al*., 1999; Wakabayashi *et al*., 2000; Ribeiro *et al*., 2002). Much of the work since the discovery of Synphilin-1 has focused on elucidating its relationships to α-synuclein aggregation, degradation, and Parkinson’s Disease. When overexpressed by itself or co-expressed with α-synuclein, Synphilin-1 forms cytosolic inclusions (O’Farrell *et al*., 2001; Xie *et al*., 2010; Lázaro *et al*., 2025). Through the use of *in vitro*, yeast, and *Drosophila* systems, several studies have indicated that Synphilin-1 inhibits α-synuclein degradation, forms cytotoxic inclusions, and induces Parkinsonian motor phenotypes (Büttner *et al*., 2010; Alvarez-Castelao and Castaño, 2011; Carvajal-Oliveros *et al*., 2021; 2023; Cao *et al*., 2025). Conversely, Synphilin-1 can also be cytoprotective in mouse and human cell systems (Tanaka *et al*., 2004; Li *et al*., 2010; Smith *et al*., 2010; Liu *et al*., 2016a; Shishido *et al*., 2019), and co-expression of Synphilin-1 and α-synuclein can promote clearance of α-synuclein and recruitment of ALS pathway components (Wong *et al*., 2008; Casadei *et al*., 2014). Beyond its interactions with α-synuclein, Synphilin-1 also plays roles in mitochondrial homeostasis, mechanotransduction, and brain development (Szargel *et al*., 2016; Kim *et al*., 2025; Savyon *et al*., 2025; Farhoud *et al*., 2026). In our current work, we provide evidence of an endogenous up-regulation of Synphilin-1 that contributes to maintaining proteostasis by modulating ubiquitinated protein dynamics and degradation.

Synphilin-1 localized around clusters of ubiquitin-conjugated proteins in both eHAP and WHAMM/JMY^DKO^ cells, suggesting that Synphilin-1 can engage these cargo and may serve the same proteostatic function in either the presence or absence of the actin nucleation factors. Consistent with previous observations that Synphilin-1 associates with the S6 regulatory ATPase of the proteasome (Marx *et al*., 2007), fluorescently-tagged Synphilin-1 often colocalized with the α3 proteasome subunit in our studies. It also colocalized with p62, which is canonically a selective autophagy receptor but can also act as a proteasomal shuttling factor and enhance UPS activity (Babu *et al*., 2005; Liu *et al*., 2016b; Fu *et al*., 2021; Lulu-Shimron *et al*., 2025). The lack endogenous Synphilin-1 staining at intense ubiquitin foci or formation of Synphilin-1 inclusions in eHAP or WHAMM/JMY^DKO^ cells strays from the exogenous expression studies, but we speculate that an adaptive up-regulation of Synphilin-1 in WHAMM/JMY^DKO^ cells may be coordinated with other factors such that the cells can maximize Synphilin-1’s ability to promote proteostasis without creating inclusion bodies.

Disruption of the actin nucleation machinery has previously been linked to defects in cell proliferation, as deletion of the Arp2/3 complex in mouse fibroblasts causes a loss of mitotic fidelity and results in genomic instability and cellular senescence (Haarer *et al*., 2023). In line with these findings, we observed that human fibroblasts lacking WHAMM and JMY and exposed to a prolonged puromycin treatment exhibited an increased frequency of micronuclei, a sign of chromosome missegregation and genomic instability. However, unlike the Arp2/3 knockout cells that senesced under genotoxic stress, the WHAMM/JMY^DKO^ cells lost viability from proteotoxic stress. The latter observations are reminiscent of the cell loss that occurred when the Arp2/3 complex knockout cells were exposed to an acute lysosomal stressor (Theodore *et al*., 2026). Since inactivation of WHAMM and/or JMY impedes the intrinsic apoptosis cascade (King *et al*., 2021; King and Campellone, 2023), the mechanisms by which WHAMM/JMY^DKO^ die remain to be determined.

Importantly, the targeted depletion of Synphilin-1 in WHAMM/JMY^DKO^ cells further exacerbated their death phenotypes and led to the buildup of p62 and ubiquitinated proteins upon exposure to puromycin. This enhanced sensitivity of WHAMM/JMY^DKO^ cells to Synphilin-1 depletion supports a model in which Synphilin-1 plays a dual role in proteostasis when autophagy is impaired: it both manipulates protein sequestration in cytosolic bodies and organizes degradation via the UPS in discrete cytosolic locations (Figure 10). Our findings add to the understanding of ALS/UPS coordination and give additional merit to the idea that protein sequestration in cytosolic structures may act as a proteotoxic waste management system for cytoprotection when the major ALS/UPS degradation pathways are altered. Determining how Synphilin-1 mediates protein dynamics and proteasomal degradation through interactions with ubiquitinated cargo, the proteasome, and other factors will likely provide further context to its physiological relevance in different cell types as well as its impact on α-synuclein during Parkinson’s Disease pathogenesis.

## MATERIALS AND METHODS

### Cell culture

Cell lines used in this study are listed in Supplemental Table S1. eHAP cells and their CRISPR-modified derivatives were described previously (King *et al*. 2021) and were cultured in Iscove’s Modified Dulbecco’s Medium (IMDM) supplemented with 10% fetal bovine serum (FBS), Glutamax, and penicillin-streptomycin. Cos7 cells (UC Berkeley cell culture facility) were cultured in Dulbecco’s Modified Eagle Medium (DMEM) supplemented with 10% FBS, Glutamax, and antibiotic-antimycotic. All cells were grown at 37°C in 5% CO_2_. Experiments were performed using cells that were in active culture for 2-10 trypsinized passages.

### Chemical treatments

For treatments with chloroquine, cells were exposed to 50μM chloroquine (Sigma Aldrich) diluted from a 50mM stock for 1-2h as noted in the Figure Legends. For treatments with puromycin, cells were exposed to media containing 0.1-5μg/mL puromycin (Sigma Aldrich) diluted from a 2.5mg/mL stock for 3h-24h as noted in the Figure Legends. For puromycin washout assays, recovery media was supplemented with 50μM chloroquine diluted from a 50mM stock or 15μM MG132 (Tocris) diluted from a 10mM stock. Equivalent volumes of media without drugs were used as controls. eHAP and WHAMM/JMY^DKO^ cell lines were also tested for their sensitivities to different concentrations of puromycin, geneticin/G418 (Fisher), and hygromycin B (Invitrogen). The minimal inhibitory concentrations were 1μg/mL puromycin, 2.0mg/mL G418, and 750μg/mL hygromycin for eHAP cells, compared to 0.5 μg/mL puromycin, 1.5 mg/mL G418, and 650 μg/mL hygromycin for WHAMM/JMY^DKO^ cells.

### DNA transfections

Plasmids were maintained in *E. coli* XL-1 Blue (Stratagene) and are listed in Supplemental Table S2. GFP-tagged human Synphilin-1 was cloned into a pEGFP vector by amplifying *SNCAIP* cDNA encoding a long isoform with a 47 amino acid extension near the N-terminal region, and a 51 amino acid extension at the C-terminus (Horizon Discovery MHS6278-202809062) by PCR with primers flanked by XhoI/NotI restriction sites. Synphilin-1 truncation mutants were generated by PCR using primers listed in Supplemental Table S2. Full length Synphilin-1 contains amino acids 1-1016, NT contains 1-396, NT-ANK1-CC contains 1-598, CC-ANK2-CT contains 562-1016, and CT contains 778-1016. mCherry-Synphilin-1 was created by subcloning the *SNCAIP* cDNA into the XhoI/NotI sites of an mCherry vector. For transient expression of GFP-Synphilin-1 and GFP-ubiquitin (Addgene #11928), Cos7 cells were transfected with 30-200ng of DNA using Lipofectamine LTX (Invitrogen) in either 24-well plates for fixation and fluorescence microscopy, or in 35mm glass-bottom dishes for live imaging. Cos7 cells stably expressing mCherry-Synphilin-1 were generated by linearizing mCherry or mCherry-Synphilin-1 plasmids with PciI and transfecting Cos7 cells with 1-2μg DNA in a 6-well plate. After 24h, cells were reseeded into 6cm dishes in media containing 1.1mg/mL G418. After 14 days, the remaining cells were trypsinized and reseeded into a 6cm dish to generate a polyclonal population.

### RNA transfections

For RNAi experiments, cells were seeded into 6-well plates and grown for 24h, transfected with 25-80nM siRNAs (Supplemental Table S2) using lipofectamine RNAiMAX (Invitrogen), reseeded into 6- or 12-well plates, or into 24-well plates containing 12mm glass coverslips, and grown for an additional 24h. Cells cultured in 6- or 12-well plates were collected and processed for immunoblotting, and cells cultured on coverslips in 24-well plates were used in fluorescence microscopy assays.

### RNA-sequencing and RT-PCR

RNA isolation, sequencing, and RT-PCRs were performed as described previously (King *et al*. 2021). eHAP, WHAMM^KO^, and WHAMM/JMY^DKO^ cells were seeded into 6cm dishes, grown for 24h, and rinsed with phosphate buffered saline (PBS) just prior to collection. RNA was isolated using TRIzol reagent (Ambion), followed by a chloroform extraction, isopropanol precipitation, and 75% ethanol wash before resuspending the RNA in water. For RNA-sequencing, RNA quality was assessed using an Agilent TapeStation 4200, and cDNA libraries were prepared based on the Illumina TruSeq Stranded mRNA sample preparation protocol. 75bp single end reads were sequenced on the Illumina NextSeq 500 to a depth of ∼30M reads each. Total reads from RNA-seq were aligned with TopHat, and differential expression was calculated with the CuffLinks/CuffDiff program. Gene expression values were given as Fragments Per Kilobase of transcript per Million mapped reads (FPKM), differential expression was calculated as the Log2(FPKM ratio KO:eHAP), and values were filtered for expression changes compared to control of at least 1.5-fold and statistically significant (q<0.05). For WHAMM^KO^, 78 genes from 13,004 total called transcripts were changed (0.6%), and for WHAMM/JMY^DKO^, 170 genes from 13,060 total transcripts were changed (1.3%). Log_2_(FPKM Ratio) values ranged from +13.07 to -13.20. For RT-PCRs, cDNA was synthesized from RNA using oligo(dT) primers (Invitrogen) and Superscript II Reverse Transcriptase (Invitrogen), then amplified using Taq polymerase (New England Biolabs) and gene-specific primers. Products were run on 1% agarose gels supplemented with ethidium bromide and visualized using a Biorad Gel Doc EZ Imager. Band intensities were quantified using the Analysis tool in LI-COR Image Studio software, and signals were normalized to GAPDH band intensities.

### Immunoblotting

For preparation of whole cell lysates to be used in analyses of LC3, GABARAP, or Synphilin-1 levels, cells were seeded into 6-well plates, grown for 24h, treated with media ± chloroquine, rinsed once with PBS, lifted with PBS containing 2mM EDTA, then centrifuged before being stored at -20°C. Pellets were resuspended in lysis buffer (20mM Tris HCl pH 7.5, 100mM NaCl, 1% TritonX-100 plus 1 mM phenylmethylsulfonyl fluoride and 10 μg/ml each aprotinin, leupeptin, pepstatin, and chymostatin), diluted in SDS–PAGE sample buffer, boiled, and centrifuged. For analyses of Synphilin-1, p62, and ubiquitinated protein levels following siRNA transfections and puromycin treatments, experiments were conducted in 12-well plates. Detached floating cells were collected by centrifugation, combined with adherent cells that had been lifted using PBS containing 1mM EDTA, and subjected to a second centrifugation. These combined cell pellets were lysed in a buffer containing 20mM HEPES pH 7.4, 50mM NaCl, 1% TritonX-100, and protease inhibitors prior to mixing with SDS-PAGE sample buffer, boiling, and centrifugation. All samples were subjected to SDS–PAGE before transfer to nitrocellulose membranes (GE Healthcare). Membranes were blocked in PBS containing 5% milk (PBS-M) before being probed with primary antibodies (Supplemental Table S3) diluted in PBS-M overnight at 4°C plus an additional 2–3h at room temperature. Membranes were rinsed twice with PBS and washed thrice with PBS + 0.5% Tween-20 (PBS-T). Membranes were then probed with secondary antibodies conjugated to IRDye-800 or IRDye-680 (Supplemental Table S3) diluted in PBS-M, rinsed with PBS, and washed with PBS-T. Blots were imaged using a LI-COR Odyssey Fc imaging system. Band intensities were determined using the Analysis tool in LI-COR Image Studio software, and quantities of proteins of interest were normalized to tubulin and/or GAPDH loading controls.

### Immunostaining

Cells grown on coverslips were fixed using 2.5% paraformaldehyde (PFA) in PBS for 30 min, washed with PBS, permeabilized with 0.1% TritonX-100 in PBS, washed, and incubated in blocking buffer (PBS containing 1% FBS, 1% bovine serum albumin (BSA), and 0.02% NaN_3_) for at least 15 min. For measuring proteasome activity, cells were treated with 500nM Me4BodipyFL-Ahx3Leu3VS probe (R&D Systems) diluted from a 250μM stock that was added to culture media 30min before fixation. For assessing cell death, cells were treated with 2μg/mL Propidium Iodide (Sigma Aldrich) diluted from a 1mg/mL stock for 30min before fixation. Cells were probed with primary antibodies (Supplemental Table S3) for 45-60min, washed, and treated with AlexaFluor-conjugated secondary antibodies, DAPI, and/or AlexaFluor-conjugated phalloidin (Supplemental Table S3) for 35min, followed by PBS washes and mounting in Prolong Gold antifade reagent (Invitrogen).

### Fluorescence microscopy

All images and videos were captured using a Nikon Eclipse Ti inverted microscope equipped with Plan Apo 100X/1.45, Plan Apo 60X/1.40, or Plan Fluor 20x/0.5 numerical aperture objectives, an Andor Clara-E camera, and a computer running NIS Elements software. Cells were viewed in multiple focal planes, and Z-series were captured in 0.2 µm steps. Images presented in the Figures represent one slice (Figure 1, Figure 7B top image, Supplemental Figures S2, S5, and S6), or three slice maximum intensity projections (Figures 2, 3, 4, 6, 7B bottom image, and 8, Supplemental Figures S3, S4, and S5). Live cell imaging was performed in a 37°C chamber (Okolab) with images captured at 10-30s intervals.

### Image processing and quantification

Image processing and analyses used ImageJ/FIJI software (Schindelin *et al*., 2012). For measurements of ubiquitin area and intensity, cells were outlined using the Freehand Selection tool and manually thresholded with the Threshold tool. Ubiquitin foci were defined as structures with clean rounded edges and intensely bright staining, whereas ubiquitin puncta were defined as small (typically less than 0.2 μm^2^), dim structures that could have clean or irregular edges. Outlines for all ubiquitin puncta/foci were generated using Analyze Particles, added to the ROI Manager, and combined to generate one region-of-interest encompassing all of the ubiquitin staining on a per cell basis. Area and intensity values were captured using the Measure tool. For pixel intensity plots, lines were drawn through puncta/foci in ubiquitin, Synphilin-1, and/or PSMA3 channels in a single Z-slice, and the Plot Profile tool was used to measure the distance and intensity along the line. For Figure 6D, the maximum intensity for ubiquitin in (ii) was set to 1, and the maximum intensity for Synphilin-1 in (i) was set to 0.5. For Figure 7C, the maximum intensity for each profile in (ii) was set to 1, and (i) was normalized to the maximum intensity in (ii) for each channel. For the number of cells, numbers of PI^+^ plus and dead/dying cells, and numbers of proteasome puncta per cell, the ImageJ Dot Counter tool was used to manually mark cell nuclei and puncta.

### Reproducibility and statistics

All conclusions were based on observations made from at least 3 separate experiments, and quantifications were based on data from 3-5 representative experiments. The sample size used for statistical tests was the number of times an experiment was performed (Lord *et al*., 2020). Statistical analyses were performed using GraphPad Prism software. Statistics for data sets comparing two conditions were determined using unpaired or paired *t* tests as noted in the Figure Legends. Statistics for data sets with 3 or more conditions were performed using ANOVAs followed by Tukey’s posthoc test unless otherwise indicated. *P* values < 0.05 were considered statistically significant.

## ACKNOWLEDGEMENTS

We thank Bo Reese at the UConn Center for Genome Innovation for guidance with RNA-seq and analyses, Adam Zweifach for discussion of statistical analyses, and Katrina Velle and Campellone Lab members for their comments on this manuscript. KGC was supported by National Institutes of Health grants GM107441, AG050774, and GM155651 (www.nih.gov). The funders had no role in study design, data collection and analysis, decision to publish, or preparation of the manuscript.

## SUPPLEMENTAL FIGURES & TABLES

**Supplemental Figure S1.**
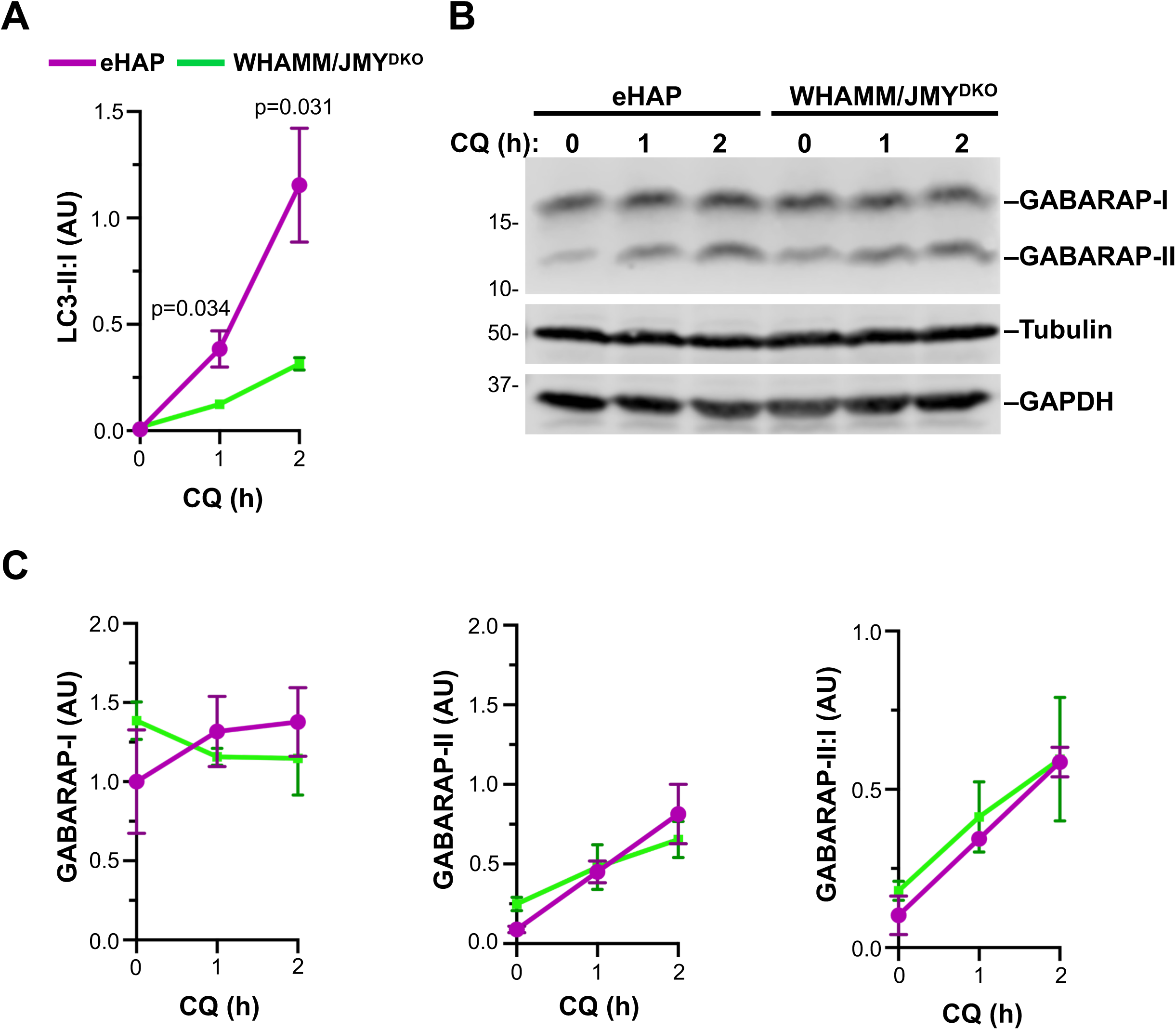
Inactivation of WHAMM and JMY reduces the LC3-II:I ratio, but does not impair GABARAP lipidation. **(A)** The ratio of LC3-II:I from Figure 1 was calculated by dividing the LC3-II band intensity by the LC3-I band intensity within each sample. Each point represents the mean from n=3 experiments ± SD. Significant *p* values are noted (unpaired *t* tests). **(B)** eHAP and WHAMM/JMY^DKO^ cells were treated with chloroquine (CQ) for 0,1, or 2h, then lysed and subjected to SDS-PAGE and immunoblotted with antibodies to GABARAP, tubulin, and GAPDH. **(C)** GABARAP-I and GABARAP-II intensities from (B) were measured relative to tubulin and GAPDH and normalized to eHAP at 0h. Each point represents the mean from n=3 experiments ± SD. The ratio of GABARAP-II:I was calculated by dividing the GABARAP-II band intensity by the GABARAP-I band intensity within each sample.

**Supplemental Figure S2.**
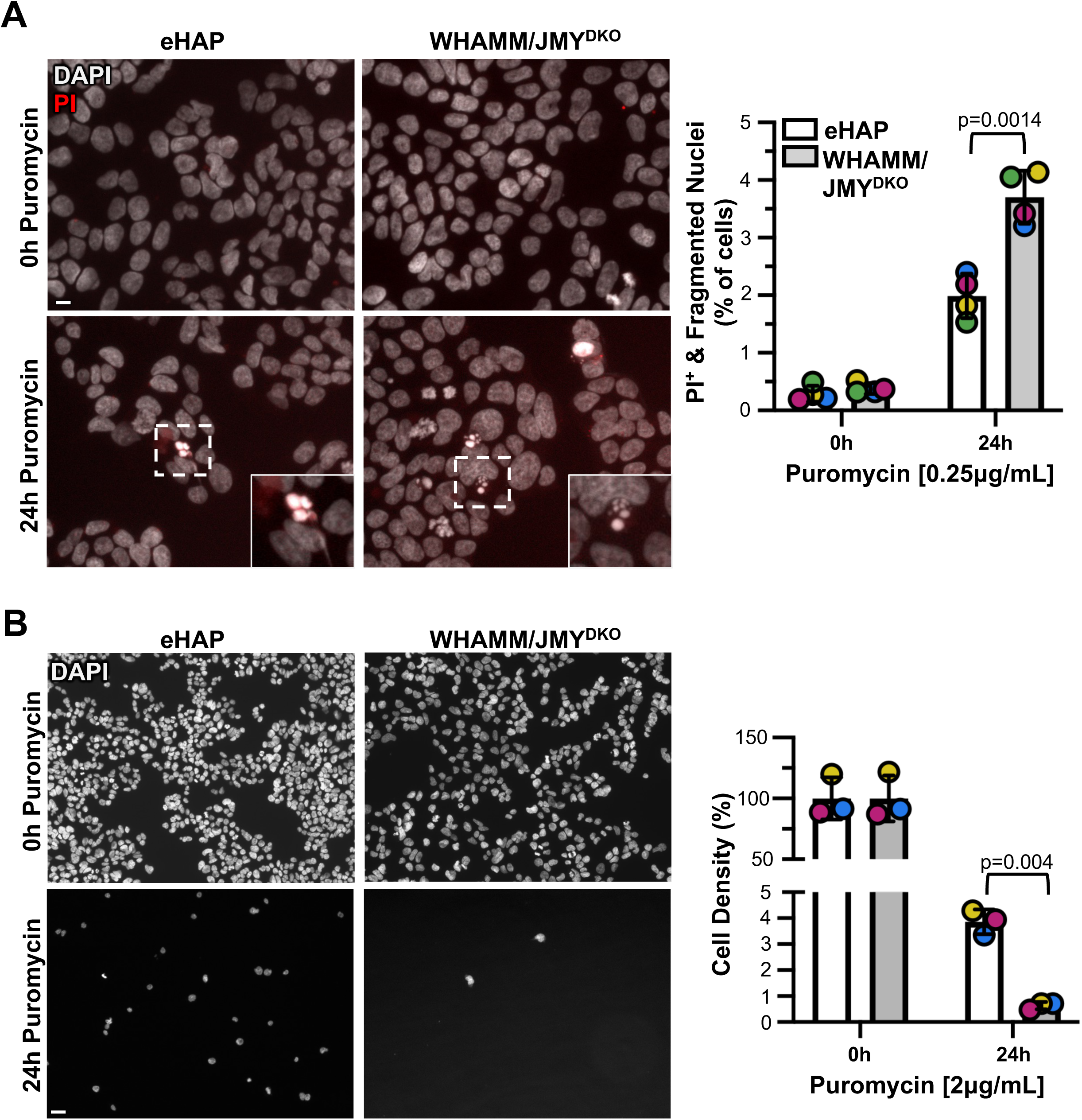
WHAMM/JMY^DKO^ cells have reduced cell viability after extended puromycin exposure. **(A)** eHAP and WHAMM/JMY^DKO^ cells were treated with 0.25μg/mL puromycin for 24h, with 2μM propidium iodide (PI, red) added to the media 30min before fixing and staining with DAPI to visualize DNA (white). Magnifications show examples of cells with condensed and fragmented DNA staining. Scale bar: 10μm. The number of cells with PI staining (PI^+^) or number of apoptotic bodies were counted and plotted as a % of the total number of cells visualized. Each symbol corresponds to the experimental average (52-798 cells counted per image with 6 images for each condition and cell line) from n=4 replicates. Each experiment is coordinated by color. Significant *p* values are noted (unpaired *t* tests). **(B)** eHAP and WHAMM/JMY^DKO^ cells were treated with 2μg/mL puromycin for 24h, then fixed and stained with DAPI. Scale bar: 20μm. The total numbers of cells in the untreated (0h) conditions were set to 100%, and the number of cells remaining after puromycin treatment were expressed as a % of the cells from the untreated condition. Each symbol corresponds to the experimental average (0-631 cells counted per image with 5 images for each condition and cell line) from n=3 replicates.

**Supplemental Figure S3.**
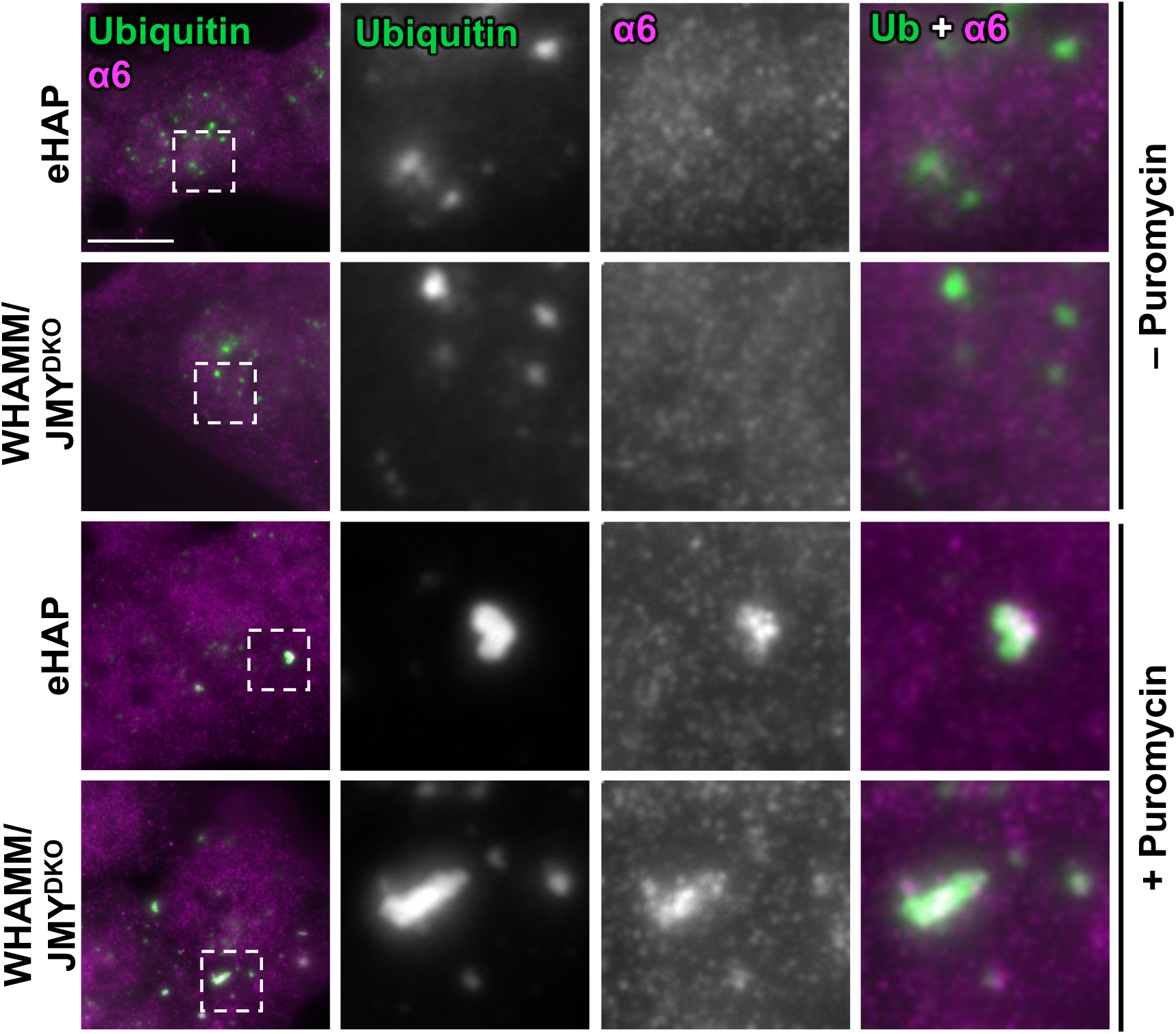
eHAP and WHAMM/JMY^DKO^ cells recruit the α6 subunit of the proteasome to ubiquitinated substrates. eHAP and WHAMM/JMY^DKO^ cells were treated with 2μg/mL puromycin for 3h and stained with antibodies to visualize ubiquitin-conjugated proteins (green) and the α6 subunit of the proteasome (magenta). Scale bar: 10μm.

**Supplemental Figure S4.**
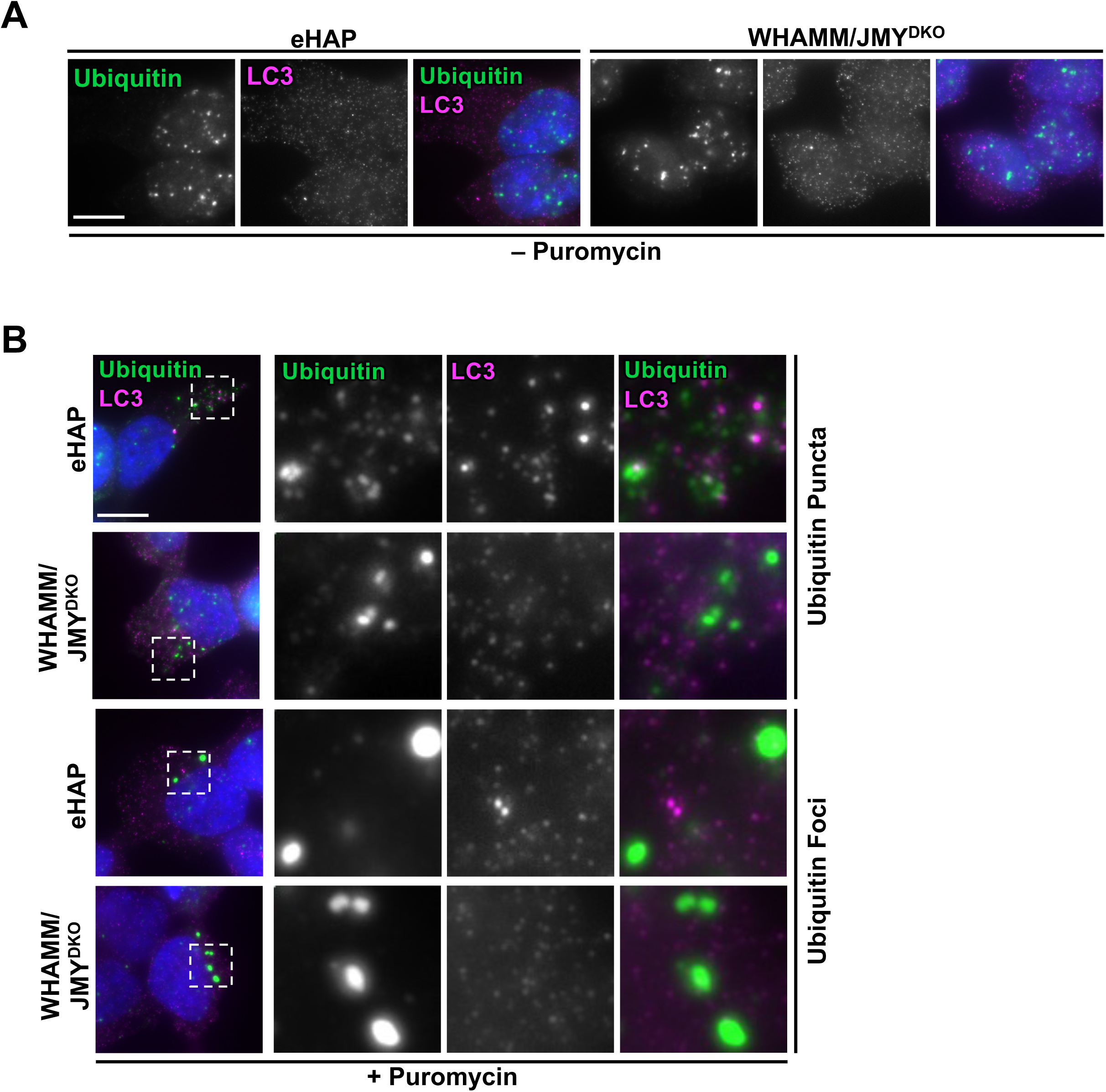
LC3 is not enriched at puromycin-induced ubiquitin foci in eHAP or WHAMM/JMY^DKO^ cells. **(A)** eHAP and WHAMM/JMY^DKO^ cells were fixed and stained with antibodies to visualize ubiquitin-conjugated proteins (green) and LC3 (magenta) and DAPI to visualize DNA. Scale bar: 10μm. **(B)** eHAP and WHAMM/JMY^DKO^ cells were treated with 2μg/mL puromycin for 3h, then fixed and stained with antibodies to visualize ubiquitin conjugated proteins (green) and LC3 (magenta). Magnifications from rows 1 and 2 highlight smaller ubiquitin puncta, while those in rows 3 and 4 highlight brighter and larger ubiquitin foci. Scale bar: 10μm.

**Supplemental Figure S5.**
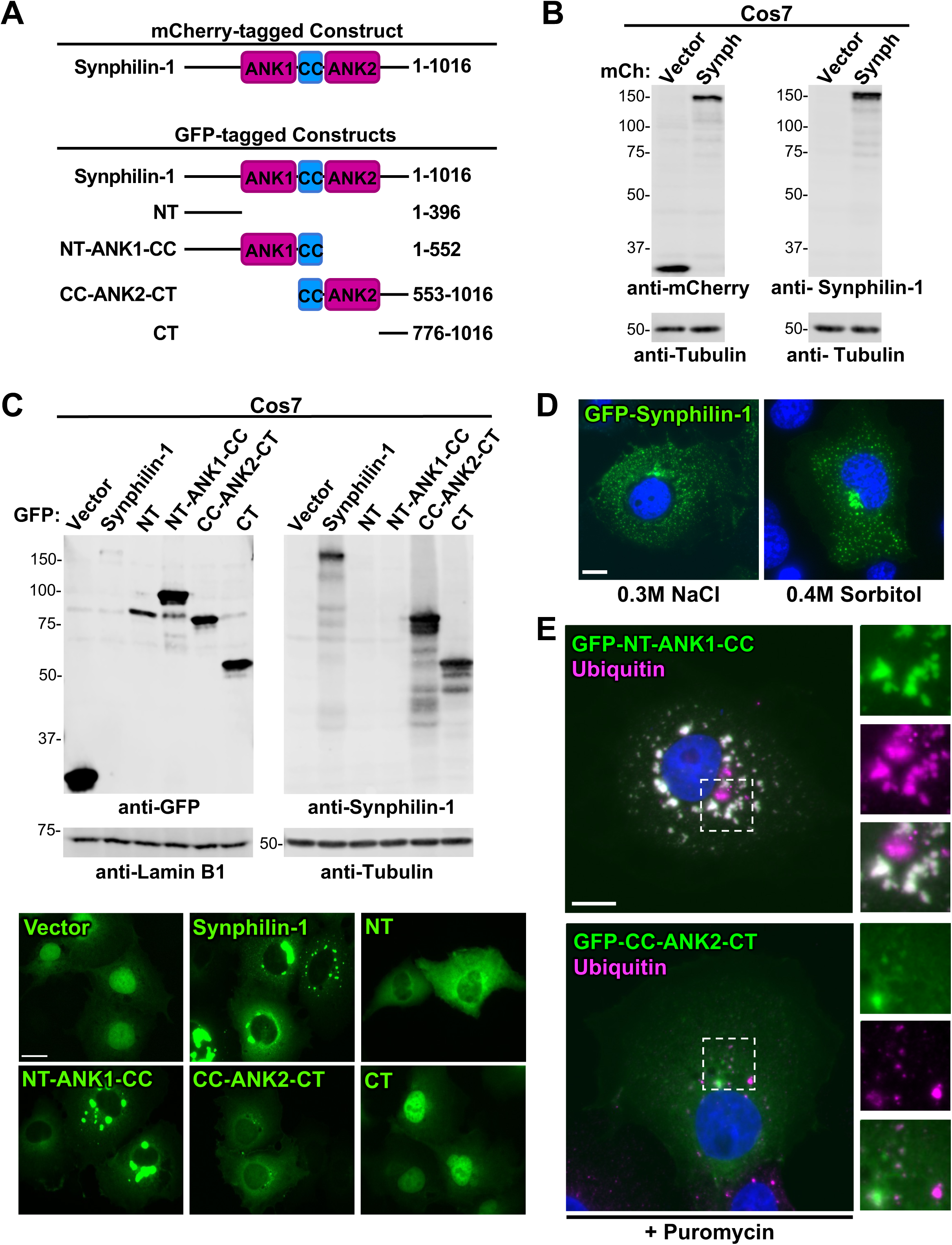
Exogenous Synphilin-1 expression drives condensate formation via its ANK1 domain, and localizes with ubiquitinated substrates and proteasomes. **(A)** Schematic of the *SNCAIP*-encoded 1016 amino acid (aa) Synphilin-1 isoform 3 protein. Subsequent truncation constructs derived from full length encompass the N-terminal domain (NT), the NT plus the ANK1 and coiled-coiled (CC) domains (NT-ANK1-CC), the CC plus the ANK2 and C-terminal domain (CC-ANK2-CT) and the C-terminal domain (CT). **(B)** Cos7 cells stably transfected with mCherry-vector or mCherry-Synphilin-1 were subjected to SDS-PAGE and immunoblotting with antibodies to mCherry, Synphilin-1, and Tubulin. **(C)** Cos7 cells were transiently transfected with GFP-vector or GFP-tagged constructs, then subjected to SDS-PAGE and immunoblotting with antibodies to GFP, Synphilin-1, Lamin B1, and Tubulin. The transfected Cos7 cells were also imaged by fluorescence microscopy. Scale bar: 20μm. **(D)** Cos7 cells transiently expressing full-length GFP-Synphilin-1 (green) were treated with 0.3M NaCl or 0.4M sorbitol for 1h, then fixed and stained with DAPI to visualize DNA (blue). Scale bar: 10μm. **(E)** Cos7 cells transiently expressing GFP-tagged NT-ANK1-CC or CC-ANK2-CT (green) were treated with 5μg/mL puromycin for 3h, then fixed and stained with antibodies to ubiquitin (magenta) and DAPI to visualize DNA (blue). Scale bar: 10μm.

**Supplemental Figure S6.**
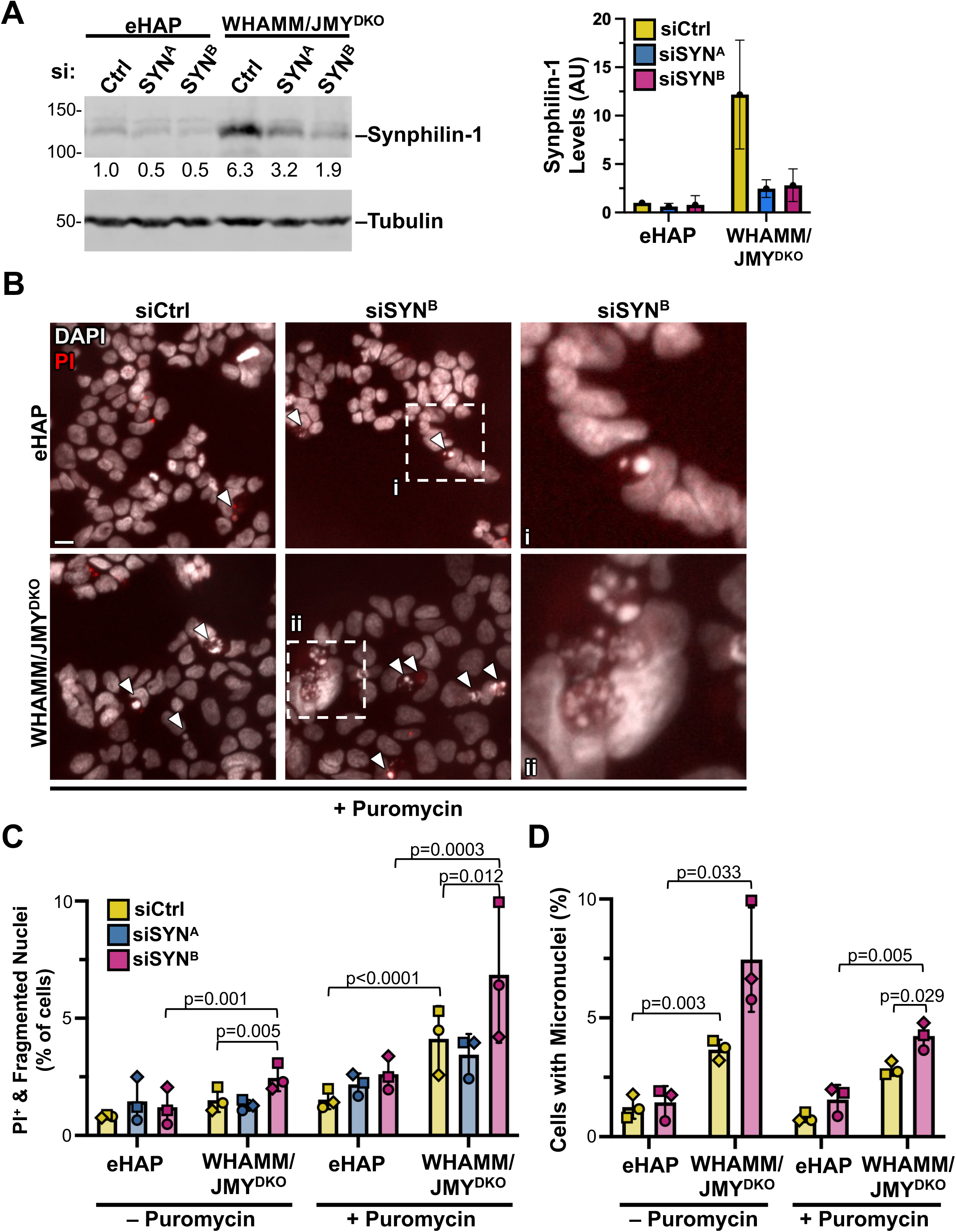
Depletion of *SNCAIP* in WHAMM/JMY^DKO^ cells reduces cell viability during proteotoxic stress. **(A)** eHAP and WHAMM/JMY^DKO^ cells were transfected with control siRNAs (siCtrl) or siRNAs targeting the *SNCAIP* transcript (siSYN) before immunoblotting with antibodies to Synphilin-1 and Tubulin. Synphilin-1 band densitometry values were normalized to Tubulin loading controls and quantified. Each bar represents the mean from n=3 experiments ± SD. **(B)** eHAP and WHAMM/JMY^DKO^ cells were transfected with siRNAs and treated with 1μg/mL puromycin for 7h with 2μM propidium iodide (PI, red) added to the media 30min before fixing and staining with DAPI to visualize DNA (white). Magnifications show examples of fragmented nuclei. Scale bar: 10μm. **(C)** The number of cells with PI staining (PI^+^) or number of fragmented nuclei were counted and expressed as a percentage of the total number of cells visualized. Each symbol corresponds to the mean from n=3 experiments ± SD (52-856 cells counted per image with 6 images for each condition and cell line). Each experiment is coordinated by symbol shape. Significant *p* values are noted (paired *t* tests for comparisons within the same cell line, and unpaired *t* tests for comparisons between cell lines). **(D)** The number of cells with at least one micronucleus was counted and divided by the total number of cells visualized. Each symbol corresponds to the mean from n=3 experiments ± SD (52-856 cells counted per image with 6 images for each condition and cell line). Significant *p* values are noted (paired *t* tests for comparisons within the same cell line, and unpaired *t* tests for comparisons between cell lines).

**Supplemental Figure S7.**
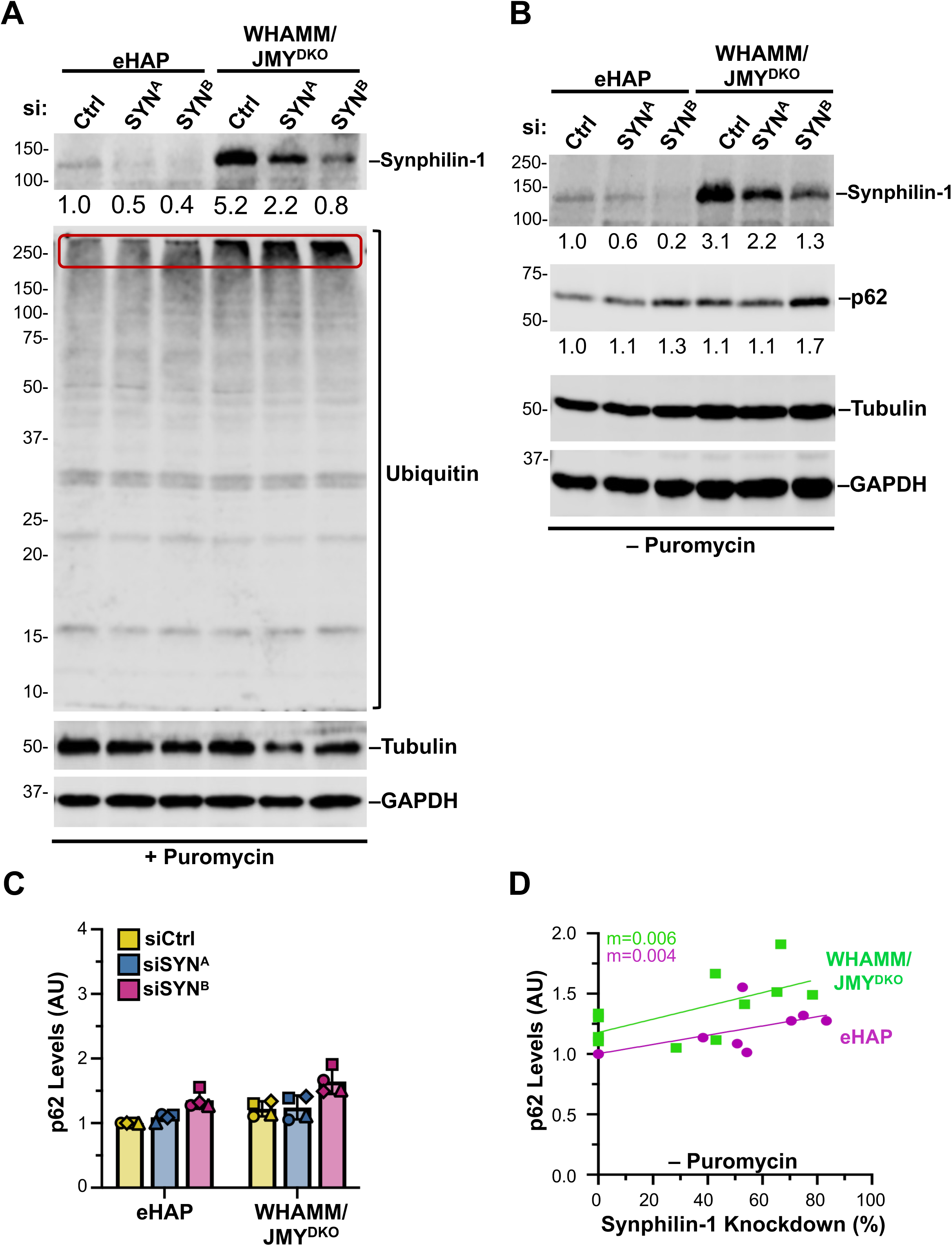
Synphilin-1 depletion in WHAMM/JMY^DKO^ cells increases the amount of ubiquitinated proteins and p62. **(A)** eHAP and WHAMM/JMY^DKO^ cells were transfected with control siRNAs or siRNAs targeting the *SNCAIP* transcript then treated with 1μg/mL puromycin for 7h before immunoblotting with antibodies to Synphilin-1, ubiquitinated proteins, Tubulin, and GAPDH. High molecular weight ubiquitin conjugates are highlighted in the red box. Densitometry values for Synphilin-1 and p62 bands were normalized to Tubulin and GAPDH controls and are shown beneath their respective blots. **(B)** eHAP and WHAMM/JMY^DKO^ cells were transfected with siRNAs before immunoblotting with antibodies to Synphilin-1, p62, GAPDH, and Tubulin. **(C)** p62 band densitometry values were normalized to Tubulin and GAPDH loading controls. Each bar represents the mean from n=4 experiments ± SD. **(D)** Densitometry values of p62 were plotted against the percentage of Synphilin-1 knockdown (calculated by dividing the densitometry value of Synphilin-1 in siSYN cells by the value of Synphilin-1 in siCtrl cells). Each point represents one sample, with samples from both siSYN^A^ and siSYN^B^ treatments shown. The slopes (m) in the linear trendline regression equations for eHAP (Y = 0.003828x + 1.003) and WHAMM/JMY^DKO^ (Y = 0.005503 + 1.179) were significantly nonzero (p=0.019 and p=0.0391 respectively).

**Table S1.**
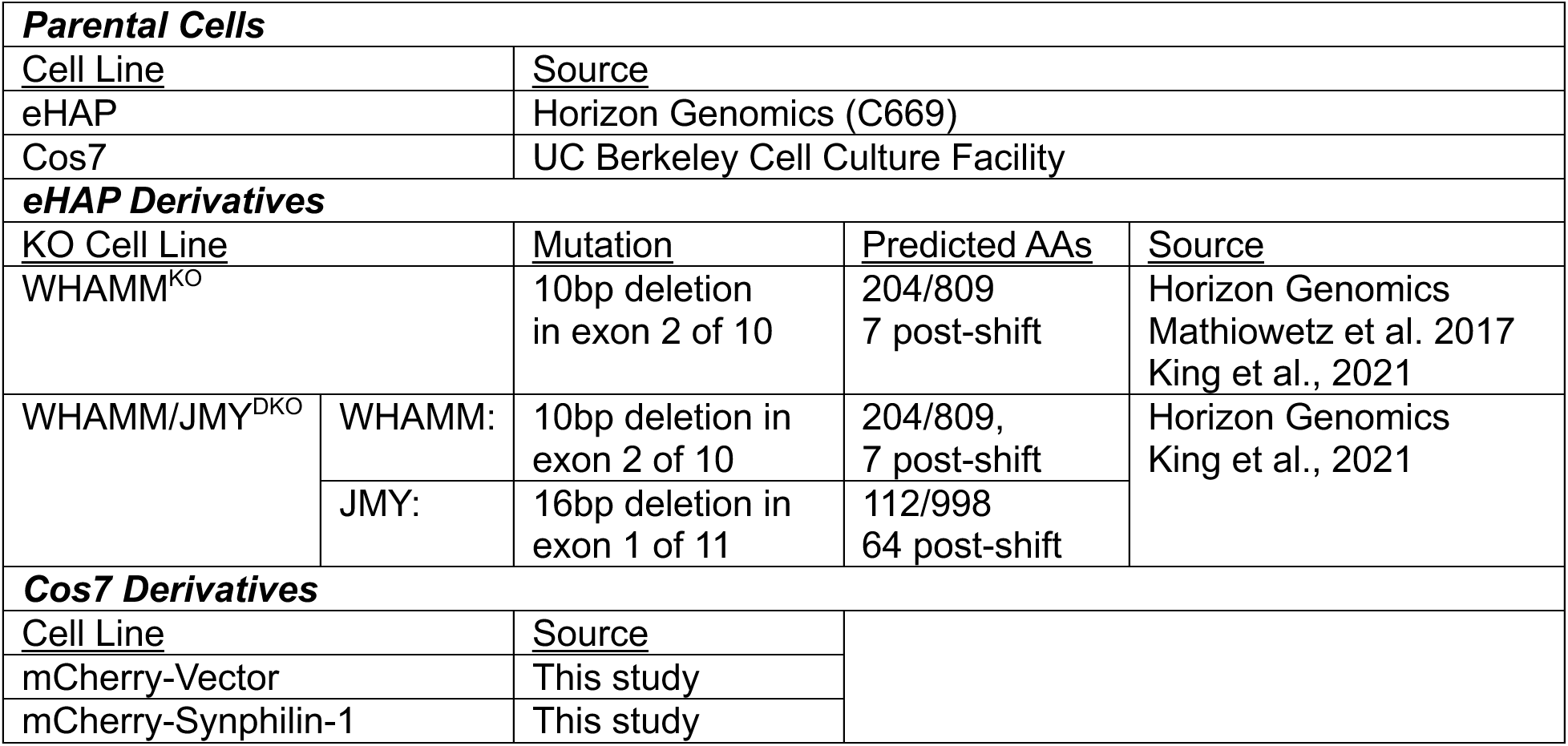
Cell Lines.

**Table S2:**
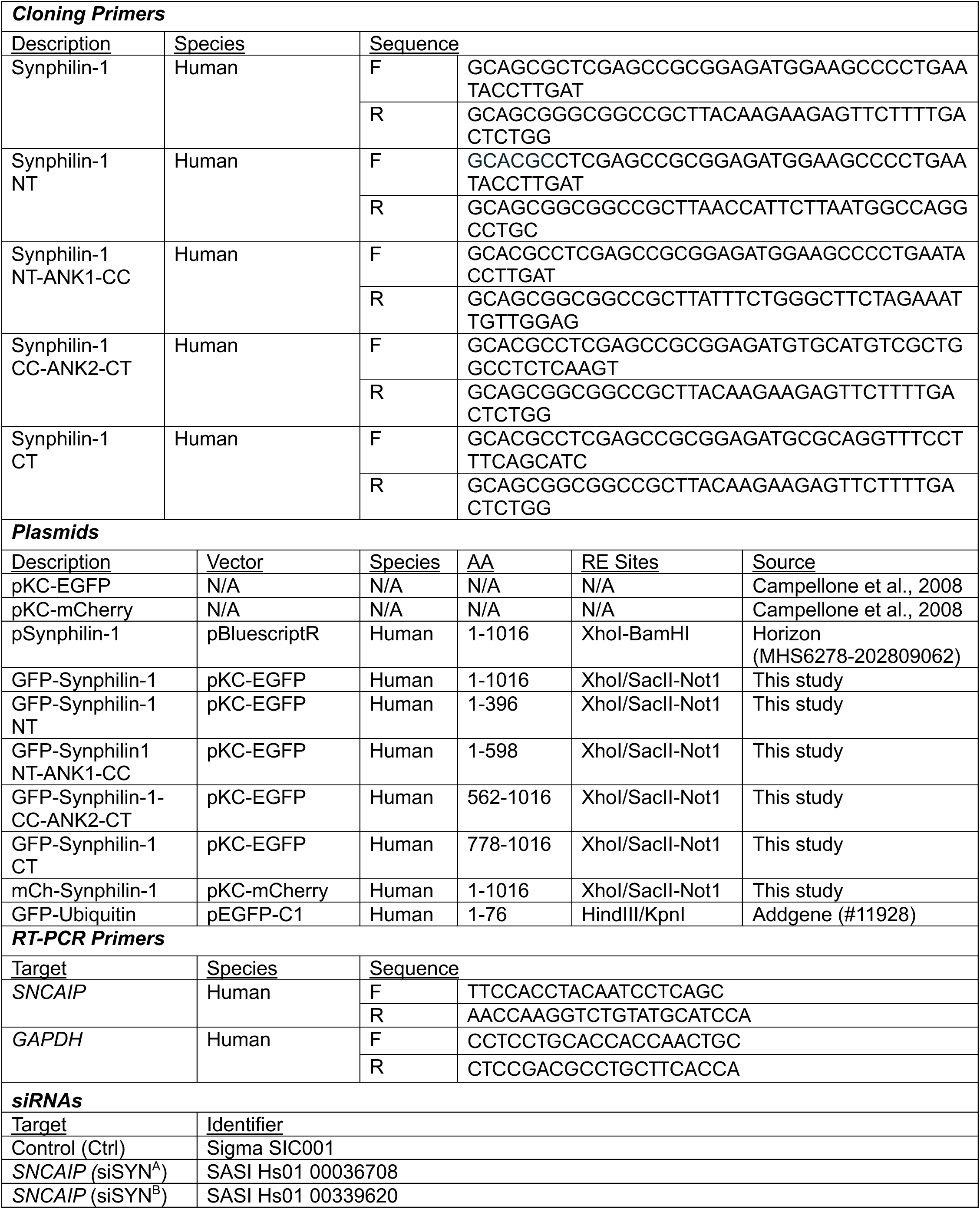
Primers, Plasmids, siRNAs.

**Table S3.**
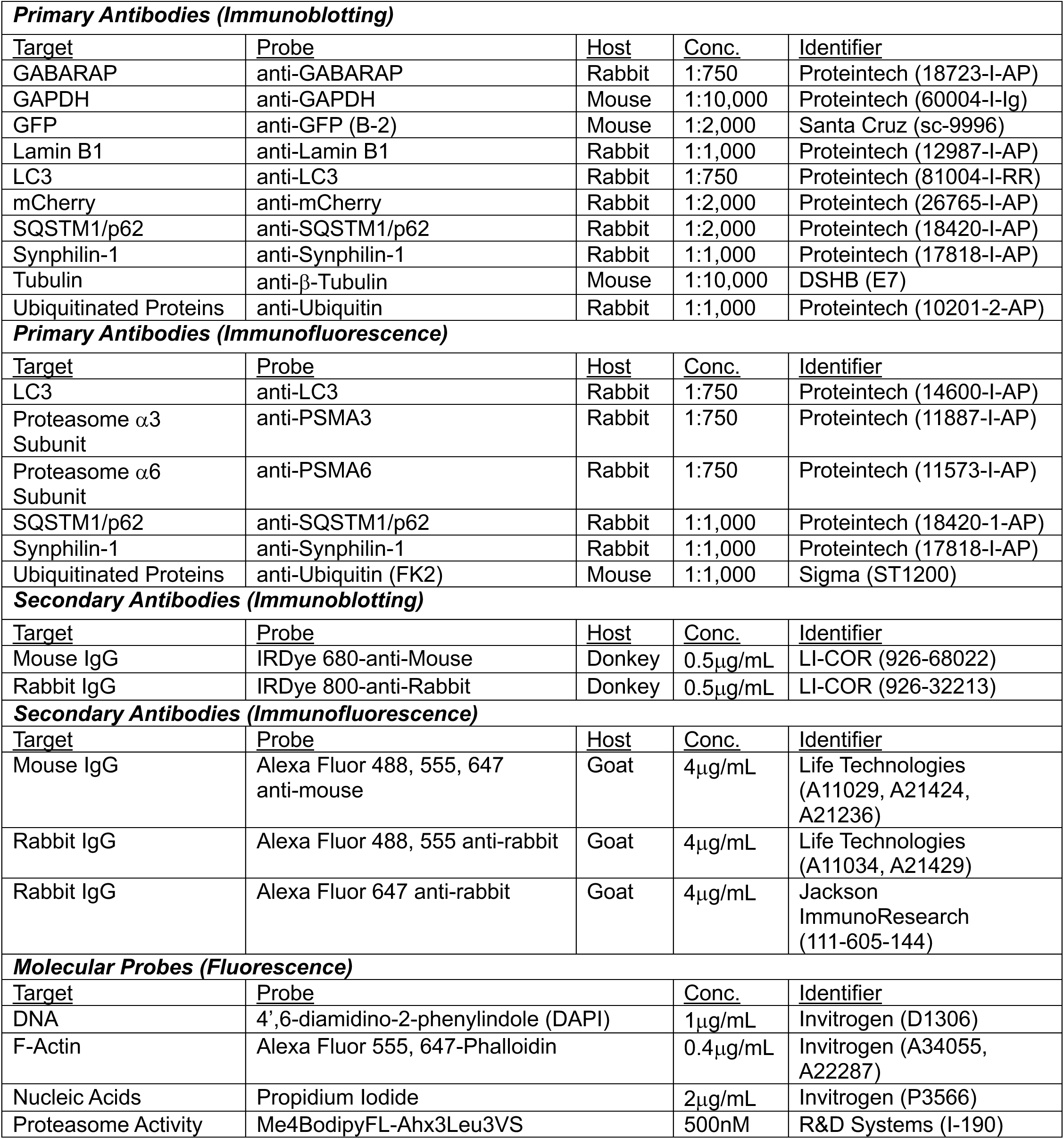
Immunoblotting and Immunofluorescence Reagents.

